# Trajectories: how functional traits influence plant growth and shade tolerance across the life-cycle

**DOI:** 10.1101/083451

**Authors:** Daniel S. Falster, Remko A. Duursma, Richard G. FitzJohn

## Abstract

Plant species differ in many functional traits that drive differences in rates of photosynthesis, biomass allocation, and tissue turnover. Yet, it remains unclear how – and even if – such traits influence whole-plant growth, with the simple linear relationships predicted by existing theory often lacking empirical support. Here we present a new theoretical framework for understanding the effect of diverse functional traits on plant growth and shade-tolerance, extending a widely-used theoretical model that links growth rate in seedlings with a single leaf trait to explicitly include influences of size, light environment, and five other prominent traits: seed mass, height at maturation, leaf mass per unit leaf area, leaf nitrogen per unit leaf area, and wood density. Based on biomass production and allocation, this framework explains why the influence of prominent traits on growth rate and shade tolerance often varies with plant size and why the impact of size on growth varies among traits. Considering growth rate in height, we find the influence of: i) leaf mass per unit leaf area is strong in small plants but weakens with size, ii) leaf nitrogen per unit leaf area does not change with size, iii) wood density is present across sizes but is strongest at intermediate sizes, iv) height at maturation strengthens with size, and v) seed mass decreases with size. Moreover, we show how traits moderate plant responses to light environment and also determine shade tolerance, supporting diverse empirical results. By disentangling the effects of plant size, light environment and traits on growth rates, our results provide a solid theoretical foundation for trait ecology and thus provide a platform for understanding growth across diverse species around the world.

## Introduction

Functional traits capture core differences in the strategies plants use to generate and invest resources (Westoby *et al.*, 2002; Wright *et al.*, 2004; Chave *et al.*, 2009). Although most woody plants have the same basic physiological function and key resource requirements (carbon, nutrients and water), species differ considerably in the rates at which resources are acquired and invested into different tissues. During the last two decades, trade-offs related to some prominent traits have been quantified, with values for traits such as leaf mass per unit leaf area, wood density, and seed size now available for up to 10% of the world’s 250000 plant species (Cornwell *et al.*, 2014). As data has accumulated, researchers are increasingly looking to traits to predict patterns in plant growth, demography, life history and performance (Poorter *et al.*, 2008; Wright *et al.*, 2010; Van Kleunen *et al.*, 2010; Adler *et al.*, 2014). In this article we outline how the growth of individual plants is influenced by various functional traits, as well as plant size and the light environment.

While the influence of traits on elements of plant physiological function has been increasingly quantified and understood, attempts at using traits to predict demographic rates have met with mixed success (Poorter & Bongers, 2006; Poorter *et al.*, 2008; Wright *et al.*, 2010; Hérault *et al.*, 2011; Paine *et al.*, 2015). In seedlings, the leaf mass per unit leaf area (LMA) – the central element of the leaf economics spectrum (Wright *et al.*, 2004) – was found to be tightly correlated with relative growth rate in plant mass (Lambers & Poorter, 1992; Cornelissen *et al.*, 1996; Wright & Westoby, 2000), as predicted from a simple mathematical model of growth rate (see below). LMA and leaf lifespan, itself closely correlated with LMA, were also linked to height growth rate for small seedlings and saplings (Reich *et al.*, 1992; Poorter & Bongers, 2006). These early successes prompted researchers to search for similar relationships in large plants. However, the results showed that in saplings and trees, LMA was not correlated with growth rate (Poorter *et al.*, 2008; Wright *et al.*, 2010; Hérault *et al.*, 2011; Paine *et al.*, 2015). Meanwhile, other traits such as wood density showed strong relationships to growth in large plants (Wright *et al.*, 2010; Hérault *et al.*, 2011), but less so in small plants (Castro-Diez *et al.*, 1998). Clearly, traits do not correlate simply and consistently to growth rate.

Recently, it has become clear that the effect of traits on plant growth can be modified by plant size (Falster *et al.*, 2011; Rüger *et al.*, 2012; Iida *et al.*, 2014; Visser *et al.*, 2016; Gibert *et al.*, 2016). A recent meta-analysis of 103 studies reporting >500 correlations by Gibert *et al.* (2016) provides the most compelling evidence. Gibert *et al.* (2016) showed that the strength of the correlation between some traits (including LMA and maximum height) and growth rate was modified by size, while for other traits (including wood density and assimilation rate per leaf area) the sign of the correlation remained the same, irrespective of size. Importantly, the direction of the correlations and their shifts with size supported predictions generated via a framework describing how traits influence growth.

Interpreting diverse empirical results seeking to link traits to growth rate is challenging because, until the paper by Gibert *et al.* (2016), we lacked any clear expectations on why the effect of traits may be moderated by size. Generating appropriate expectations is one of the primary roles for theory (Kokko, 2007). A widely-used equation for seedlings suggests that, all else being equal, a seedling’s relative growth rate in mass is linearly and negatively related to LMA (Lambers & Poorter, 1992; Cornelissen *et al.*, 1996; Wright & Westoby, 2000). An extension of the model suggests a similar relationship should hold at larger plant sizes (Enquist *et al.*, 2007). But as noted above, the prediction for large plants is not supported by empirical results. Meanwhile, theoretical predictions on how other traits should influence growth are largely absent. Without further guidance, many researchers expected traits to linearly map onto growth rates, i.e. there is a “fast” and “slow” growth end for each trait (e.g. Grime, 1977; Poorter *et al.*, 2008; Chave *et al.*, 2009; Paine *et al.*, 2015).

One possible reason that theoretical predictions have either been lacking or not been supported is that the effects of traits in existing theory are realised mainly via influences on primary productivity (photosynthesis - respiration) (Wright & Westoby, 2000; Enquist *et al.*, 2007). By contrast, the physiology of traits such as LMA and wood density suggests they have influence by altering the allocation of biomass among different tissues, rather than changing mass production (Falster *et al.*, 2011; Duursma & Falster, 2016; Gibert *et al.*, 2016). A second concern for theory focussed on mass production is that measuring mass production is really only practical for small plants that can be easily harvested. For larger plants, growth is more often measured as increment in either diameter or height. For example, stem diameter growth has been measured for many decades across many thousands of plots across the globe (Purves & Pacala, 2008; Anderson-Teixeira *et al.*, 2015; Kunstler *et al.*, 2016). Theory considering only primary production therefore offers limited insight.

Here we show how a new mechanistic growth model – called plant (Falster *et al.*, 2016) – can explain diverse empirical phenomena, including a size-dependent effect of traits on growth and an effect of traits on shade tolerance (Table 1), and thereby offer new insights into the way traits influence plant demography across the life-cycle. Broadly, the plant model builds on several approaches to modelling production and allocation of biomass (e.g. Givnish, 1988; Yokozawa & Hara, 1995; Mäkelä, 1997; Moorcroft *et al.*, 2001; Sitch *et al.*, 2008; Falster *et al.*, 2011; King, 2011). Our primary focus in this article is to explain the pattern that has been gradually emerging – that the effect of traits on plant growth is modified by plant size (Rüger *et al.*, 2012; Iida *et al.*, 2014; Gibert *et al.*, 2016). Based on the same decomposition of growth rates as is implemented in the plant model and used below (from Falster *et al.*, 2011), Gibert *et al.* (2016) argued conceptually why the effect of traits on growth should change with size. Here we extend the results of Gibert *et al.* (2016) to provide a full, functioning model and use this to show, from the point of view of primary production and allocation, how and why the effect of some traits on growth rates changes with size. We also show how our approach can account for other phenomena, including changes in growth and shade tolerance with traits, individual size, and light environment. Our view is that trait-based approaches – which aim to explain differences among species – should be integrated within a general model of plant growth, and thus should also be able to capture patterns of growth through ontogeny. Growth rates tend to show hump-shaped relationship with size, when expressed as either height (Sillett *et al.*, 2010; King, 2011) or diameter growth (Canham *et al.*, 2004, 2006; Hérault *et al.*, 2011) or biomass production (Givnish, 1988; Koch *et al.*, 2004). In contrast, the growth rate of standing plant mass continues to increase with size (Sillett *et al.*, 2010; Stephenson *et al.*, 2014). All growth measures decrease sharply with size when expressed as relative growth rates (Rees *et al.*, 2010; Iida *et al.*, 2014). Shade tolerance also varies among species, correlates with traits (Messier *et al.*, 1999; Lusk *et al.*, 2008; Poorter & Bongers, 2006), and tends to decrease with increasing size (Givnish, 1988; Kneeshaw *et al.*, 2006; Lusk *et al.*, 2008). These diverse phenomena – summarised in Table 1 – are deserving of a comprehensive and integrated explanation.

**Table 1:** Key empirical phenomena explained via the framework presented in this paper. Select references are provided for each phenomena, with preference for meta-analyses, where available. The traits considered are: seed mass (*ω*), height at maturation (*H*_mat_), leaf nitrogen content per unit leaf area (*ν*)^1^, leaf mass per area (*φ*)^2^, and wood density (*ρ*). See Tables 2 and 3 for further details. Abbreviations: rgr=relative growth ratel LAI = leaf area index.

| Phenomena | References |
| --- | --- |
| **Change in growth rate with increasing size (Fig. 1)** |  |
| Net biomass production: Hump-shaped | Givnish (1988); Koch *et al.* (2004) |
| Plant mass: Increasing | Sillett *et al.* (2010); Stephenson *et al.* (2014) |
| Height: Hump-shaped | Ryan *et al.* (2006); Sillett *et al.* (2010); King (2011) |
| Stem-diameter: Hump-shaped | Canham *et al.* (2004, 2006); Hérault *et al.* (2011) |
| Stem-area: unknown |  |
| RGR (all variables): decreasing | Rees *et al.* (2010); Iida *et al.* (2014) |
| **Effect of traits on growth rate (Fig. 4)** |  |
| $\omega$: low values produce smaller seedlings, resulting in lower absolute growth but higher RGR. | Meta-analysis by Gibert *et al.* (2016) |
| $H_{\text{m}}$: low values slow growth, but only at larger sizes | Meta-analysis by Gibert *et al.* (2016) |
| $\nu$: low values increase growth irrespective of size, but only in high light | Meta-analysis by Gibert *et al.* (2016) |
| $\phi$: low values increase growth when small, not at mid-large sizes | Meta-analysis by Gibert *et al.* (2016) |
| $\rho$: low values increase growth, except at largest sizes | Meta-analysis by Gibert *et al.* (2016) |
| **Responsiveness of growth rate to changes in light, $E$ (Fig. 4)** |  |
| $\nu$: high values respond more to $E$, changes ranking of growth rate across species | |
| $\phi$: low values respond more to $E$, does not change ranking of growth rate across species | |
| $\rho$: low values respond more to $E$, does not change ranking of growth rate across species | Rüger *et al.* (2012) |
| **Shade tolerance, wplcp (Fig. 5)** |  |
| Decreasing with size | Givnish (1988); Kneeshaw *et al.* (2006); Lusk *et al.* (2008) |
| $\nu$: low values more shade tolerant | Messier *et al.* (1999); Craine & Reich (2005); Baltzer & Thomas (2007) |
| $\phi$: low values less shade tolerant | Messier *et al.* (1999); Poorter & Bongers (2006); Baltzer & Thomas (2007); Lusk *et al.* (2008)<sup>3</sup> |
| → stands have higher LAI | Reich *et al.* (1992); Gower *et al.* (1993); Niinemets (2010) |
| $\rho$: low values less shade tolerant | Osunkoya (1996) |

<sup>1</sup> Similar responses are predicted for maximum photosynthetic rate per leaf area, or dark respiration rate per leaf area. Here these terms are directly related to $\nu$ .
<sup>2</sup> Similar responses are predicted for leaf lifespan. In our analysis this terms is directly related to $\phi$ .
<sup>3</sup> Note also that the article by Baltzer & Thomas (2007) reports a very strong relationship between WPLCP and respiration rate measured on either area or mass basis.

## Framework for understanding the effects of traits on growth

The plant model builds on the widespread approach used in many vegetation models of explicitly modelling the amounts of biomass in different tissues within a plant (e.g. Givnish, 1988; Mäkelä, 1997; Moorcroft *et al.*, 2001; Sitch *et al.*, 2008; Falster *et al.*, 2011; King, 2011; De Kauwe *et al.*, 2014) (Fig. 1a). We consider the masses *M*_*i*_, areas *A*_*i*_, and diameters *D*_*i*_ of tissues, where the subscripts indicates tissue type: l=leaf, b=bark and phloem, s=sapwood, h=heartwood, r=root, a=alive (l+b+s+r), t=total (l+b+s+h+r), and st=stem total (s+b+h). The total mass of living tissue is then *M*_a_ = *M*_l_ + *M*_b_ + *M*_s_ + *M*_r_, and the standing mass of the plant is *M*_t_ = *M*_a_ + *M*_h_. A summary of all variables, units and definitions is given in Table 2–3, with further details on the parameter values applied given in Tables S1-S2.

**Figure 1:**
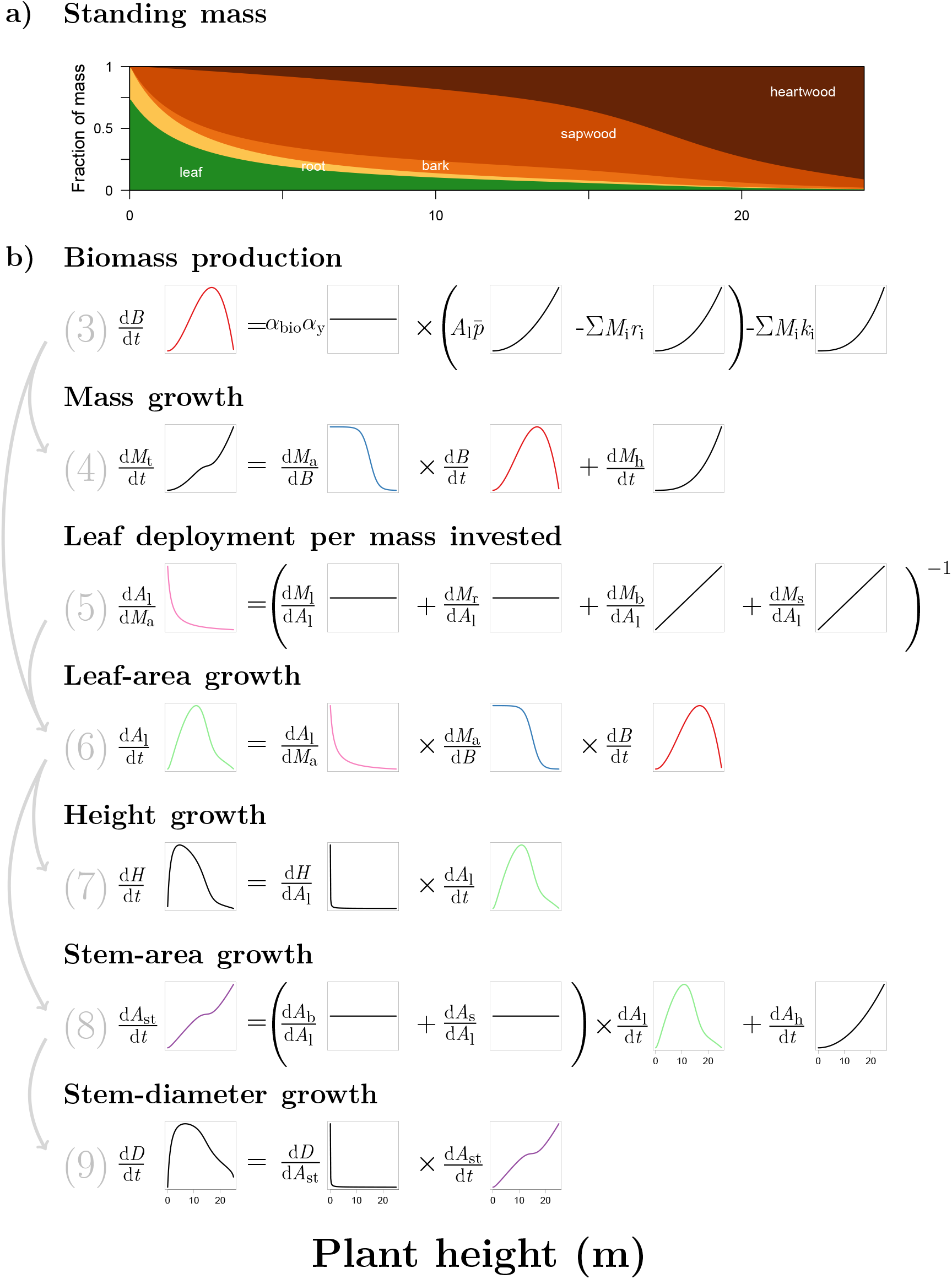
Conceptual framework linking growth rate to plant size and traits. **a)** Shows how the distribution of mass in a typical plant varies with size. **b)** Equations describing the rates of biomass production and growth in various dimensions of the plant. In the first line the symbol Σ means “sum”, across tissues, where *i* = l, b, sr. The grey numbers indicate equation numbers referred to in the main text. The insets show how the different metrics change intrinsically with plant height, when applying the “functional-balance” model in Table 4. Colours highlight where the same metric appears repeatedly in different equations. For a full list of variable names see Table 2.

**Table 2:** Key variables of the FF16 physiological model from plant, and used in this analysis. For mass (*M*), respiration (*r*), and turnover (*k*) variables, subscripts refer to any of the following tissues: l = leaves, b = bark, s = sapwood, r = roots, a = all living tissue. For area *A* variables, subscripts refer to any of the following: l = leaves, st = total stem cross-section, s = sapwood cross-section, b = bark cross-section, h = heartwood cross-section.

| Symbol | Unit | Description |
| --- | --- | --- |
| **Environmental variables** |  |  |
| $E$ | | Canopy openness |
| **State variables (may vary through ontogeny)** |  |  |
| $H$ | m | Height of a plant |
| $B$ | kg | Biomass originating from parent plant |
| $D$ | m | Stem diameter |
| $M_i$ | kg | Mass of tissue type $i$ retained on plant |
| $A_i$ | m<sup>2</sup> | Surface area or area of cross-section of tissue type $i$ |
| $p, \bar{p}$ | mol yr<sup>-1</sup> m<sup>-2</sup> | Photosynthetic rate per unit area |
| $r_i$ | mol yr<sup>-1</sup> kg<sup>-1</sup> | Respiration rate per unit mass of tissue type $i$ |
| $k_i$ | yr<sup>-1</sup> | Turnover rate for tissue type $i$ |
| **Traits (constant through ontogeny)** |  |  |
| $\omega$ | kg | Seed mass |
| $H_{\text{mat}}$ | m | Height at maturation |
| $\nu$ | kg m<sup>-2</sup> | Leaf nitrogen per unit leaf area |
| $\phi$ | kg m<sup>-2</sup> | Leaf mass per area |
| $\rho$ | kg m<sup>-3</sup> | Wood density |
| **Other parameters (constant through ontogeny)** |  |  |
| $\alpha_y$ | | Yield – fraction of carbon fixed converted into mass |
| $\alpha_{\text{bio}}$ | kg mol<sup>-1</sup> | Biomass per mol carbon |
| $\eta_c$ | | Crown-shape parameter |
| $\theta$ | | Sapwood area per unit leaf area |
| $\alpha_{l1}$ | m | Height of plant with leaf area of 1m<sup>2</sup> |
| $\alpha_{r1}$ | kg m<sup>-2</sup> | Root mass per unit leaf area |
| $\alpha_{b1}$ | | Ratio of bark area to sapwood area |

**Table 3:** Key trade-offs for the five traits considered here. For each trait we list the benefit and cost of increased trait values, as encoded into the plant model.

| Name | Benefit | Cost | References |
| --- | --- | --- | --- |
| Seed mass, $\omega$ | ↑ size seedlings | ↓ maternal fecundity | Moles & Westoby (2006) |
| Height at maturation, $H_{\text{mat}}$ | ↑ growth | ↓ fecundity | Thomas (2011); Wenk & Falster (2015) |
| Leaf nitrogen per unit leaf area, $\nu$ | ↑ photosynthesis in high light | ↑ respiration rate | Wright *et al.* (2004) |
| Leaf mass per unit leaf area, $\phi$ | ↓ leaf turnover | ↑ cost building leaf | Wright *et al.* (2004) |
| Wood density, $\rho$ | ↓ sapwood turnover | ↑ cost building stem | Chave *et al.* (2009) |

We assume growth is fundamentally driven by biomass production and its subsequent distribution throughout the plant. Applying a standard approach, the amount of biomass available for growth, 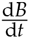 is given by the difference between income (photosynthesis) and losses (respiration and turnover) within the plant (Mäkelä, 1997; Thornley & Cannell, 2000):

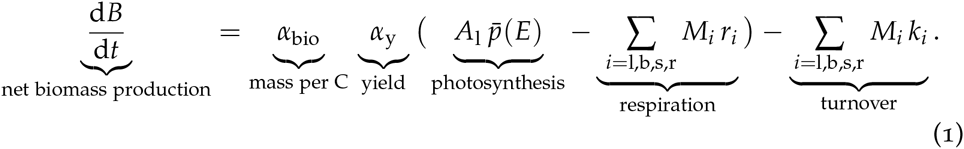

Photosynthesis is the product of the average photosynthetic rate per unit leaf area 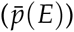 and total leaf area (*A*_l_). We assume that 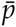 increases with canopy openness *E*, as per a standard light-response curve (Fig. 2; for details, see Supplementary Materials), and respiration and turnover rates of different tissues are constants that might differ with traits (see below). The constants *α*_y_ and *α*_bio_ account for growth respiration and the conversion of CO_2_ into units of biomass, respectively. While the plant model can easily accommodate competitive shading via influences on *E*, in this analysis we grow individual plants under a fixed light environment, so that we can better understand the intrinsic trait- and size-related effects. We also do not yet consider any traits influencing levels of self-shading. Also, many vegetation models use a more-detailed physiological model for calculating 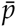 and *r*_i_, e.g. as functions of temperature. Such detail is not needed here because it will not change general model behaviour, even it would alter he absolute values of predicted growth rates. Even with detailed models, 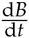 always comes down to simple subtraction of income and losses.

**Figure 2:**
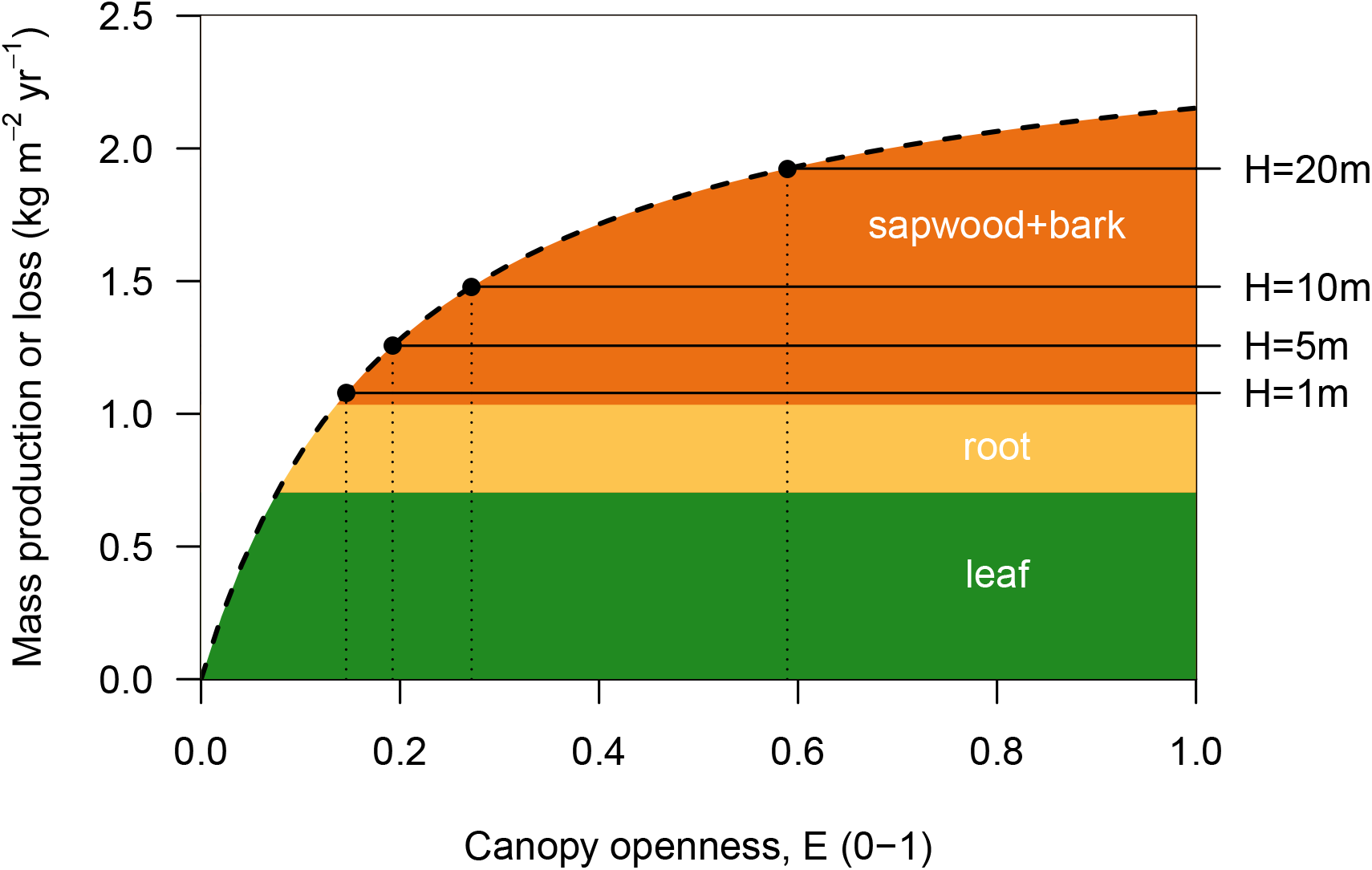
Conceptual framework linking shade-tolerance to plant size and traits, adapted from Givnish (1988). Shade tolerance is the level of canopy openness *E* where photosynthetic income (dashed line) intersects with the sum of respiration and turnover costs over tissues (solid black lines). The black circles indicate the point of intersection for plants with different heights. All income and cost is expressed in units of dry mass produced per unit leaf area per year. Note the costs of sapwood and bark increases with height. Traits can impact on shade tolerance by, in the case of leaf nitrogen content, shifting the income line up or down, or in the case of leaf mass per unit leaf area or wood density, causing the cost components to increase or decrease.

### Classic model for mass-based relative growth rate

Earlier studies focussing on seedlings used a mathematical model to relate the trait leaf mass per area (*φ*) to relative growth rate in mass (Blackman, 1919; Lambers & Poorter, 1992; Cornelissen *et al.*, 1996; Wright & Westoby, 2000). To aid comparison with this literature, we will first show how that model is derived from eqn 1. In fact, the seedling model is a special case of the more-extended approach described in the following sections and can be derived from eqn 1 as follows. For seedlings, which are young and mostly consist of leaf biomass, we can as a first approximation ignore all turnover terms as well as the respiration terms for non-leaf tissues in eqn 1. Net production (from eqn 1) then becomes a linear function of leaf area, making relative growth rate in mass a linear function of *φ*:

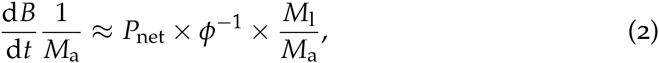

where 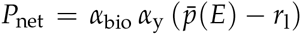. Although eqn 2 captures patterns of growth in seedlings in relation to *φ* (Wright & Westoby, 2000), this approximation does not easily extend to the variables that are routinely collected for large trees: namely plant height (*H*), stem-basal area (*A*_st_) or stem diameter *D*. The derivations below makes these links clear.

### Decomposition of growth rates into components

To model growth in either plant height (*H*), leaf area (*A*_l_), basal stem area (*A*_st_), standing mass (*M*_t_) or stem diameter (*D*) requires that we account not just for net mass production, but also for the costs of building new tissues, allocation to reproduction, and architectural layout. Mathematically, these growth rates can be decomposed into a product of physiologically relevant terms (Falster *et al.*, 2011; Gibert *et al.*, 2016). Relevant equations are given in Fig. 1b, (eqns 4-9). As is evident in eqn Fig. 1b, the growth rate in plant weight (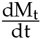; eqn 4), leaf area (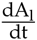; eqn 6), height (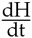; eqn 7), stem basal area (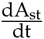; eqn 8), and stem diameter (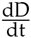; eqn 9) all share some terms in common. Many of the terms in Fig. 1 also vary intrinsically with size. The insets in Fig. 1b show the size-related patterns in different variables for a typical plant, obtained from applying the specific allocation model in the next section.

The growth rate of all size metrics in eqns 4-9 depends on the product of biomass production 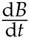 (from eqn 1) and the fraction of biomass allocated to growth of the plant, 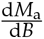, which varies from 0-1. The remaining 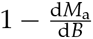 fraction of mass produced is allocated to reproduction. In plants, 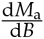 starts high, 1.0 for seedlings, and then decreases through ontogeny, potentially to zero in fully mature plants (Wenk & Falster, 2015). Note also that 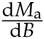 is the allocation of biomass *after* replacing parts lost due to turnover. So a plant with 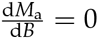 will continue to produce some new leaves and increase in stem diameter, even if the net amount of live mass *M*_a_ is not increasing.

The growth rate in the total standing mass of the plant (eqn 4) is then the sum of heartwood formation (=sapwood turnover) and any increment in live mass.

The remaining growth rates (eqns 6-9) all depend on another variable, 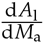, that accounts for the marginal cost of deploying an additional unit of leaf area, including construction of the leaf itself and supporting bark, sapwood and roots (eqn 5). The inverse of this term, 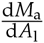, is the whole plant construction cost per unit leaf area, which can be further decomposed as a sum of tissue-level construction costs per unit of leaf area, with one of these being 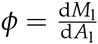: the trait leaf mass per area.

The rate of height growth (eqn 7) depends on an additional term, 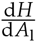: the growth in plant height per unit growth in leaf area. This variable accounts for the architectural strategy of the plant. Some species tend to leaf out more than grow tall, while other species emphasise vertical extension (Poorter & Bongers, 2006).

The rate of stem-basal-area growth (eqn 8) can be expressed as the sum of increments in sapwood, bark and heartwood areas (*A*_s_, *A*_b_, *A*_h_ respectively): 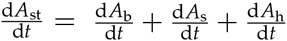. These in turn are related to ratios of sapwood and bark area per leaf area, and sapwood turnover (see eqn 8).

Finally, the rate of stem-diameter growth (eqn 9) is given by a geometric relationship between stem diameter (*D*) and stem area (*A*_st_). Note that we make no assumptions about the relationship of stem diameter to height or leaf area: these arise as emergent properties, via integration of stem turnover (eqns 13-22).

### Shade tolerance

Eqn 1 can also be used to estimate a measure of shade tolerance: the light level at which a plant’s photosynthetic gains just balance the costs of tissue turnover and respiration (Givnish, 1988; Baltzer & Thomas, 2007; Lusk & Jorgensen, 2013) (the “whole-plant-light-compensation point”, WPLCP). In general, average assimilation rate per leaf area 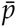 increases with light level or canopy openness, *E*. The WPLCP can be estimated by solving for the the value of canopy openness *E* = *E*\* giving 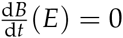 (Fig. 2). From eqn 1, this occurs when

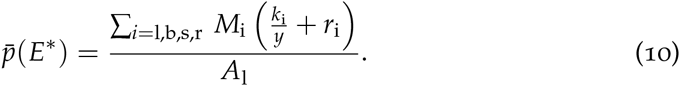

The WPLCP occurs at the points where the photosynthetic production (per unit leaf area) line intersects with the sum of maintenance and respiration costs (per unit leaf area) for each tissue (Fig. 2). Traits can influence the WPLCP if they effect either carbon uptake or costs (respiration, turnover). Also, since the amount of stem support increases with plant height, the WPLCP also naturally increases with height (Givnish, 1988) (Fig. 2).

### A functional-balance model for plant construction

It worth noting that because eqns 4–10 are derived using standard rules of addition, multiplication and differentiation, they hold for any potential growth model where biomass allocation is important. To make explicit predictions via this framework, then requires an explicit model of plant construction and function; i.e. we must put quantify all the terms in Fig. 1b.

The plant package adopts a model of plant construction and function that can be considered a first-order functional-balance or function-equilibrium model, similar to those implemented by Mäkelä (1997) and Moorcroft *et al.* (2001). We could also call it “isometric”, because the assumptions see area-based metrics scaling to the first-power of other area-based metrics, and to the square-power of length-based metrics, such as height (Huxley, 1932). Table 4 provides key equations of the model (see Falster *et al.* 2016 for full derivation). In particular, we assume that as a plant grows:

**Assumption1:** Its height scales to the 0.5 power of its leaf area (eqn 13).
**Assumptions2:** The cross-sectional area of sapwood in the stem is proportional to its leaf area (eqn 14).
**Assumptions3:** The cross-sectional area of bark and phloem in the stem is proportional to its leaf area (eqn 15).
**Assumptions4:** The cross-sectional area of root surface area and therefore mass is proportional to its leaf area (eqn 16).
**Assumption5:** The vertical distribution of leaf within the plant’s canopy, relative to the plant’s height, remains constant.

**Table 4:**
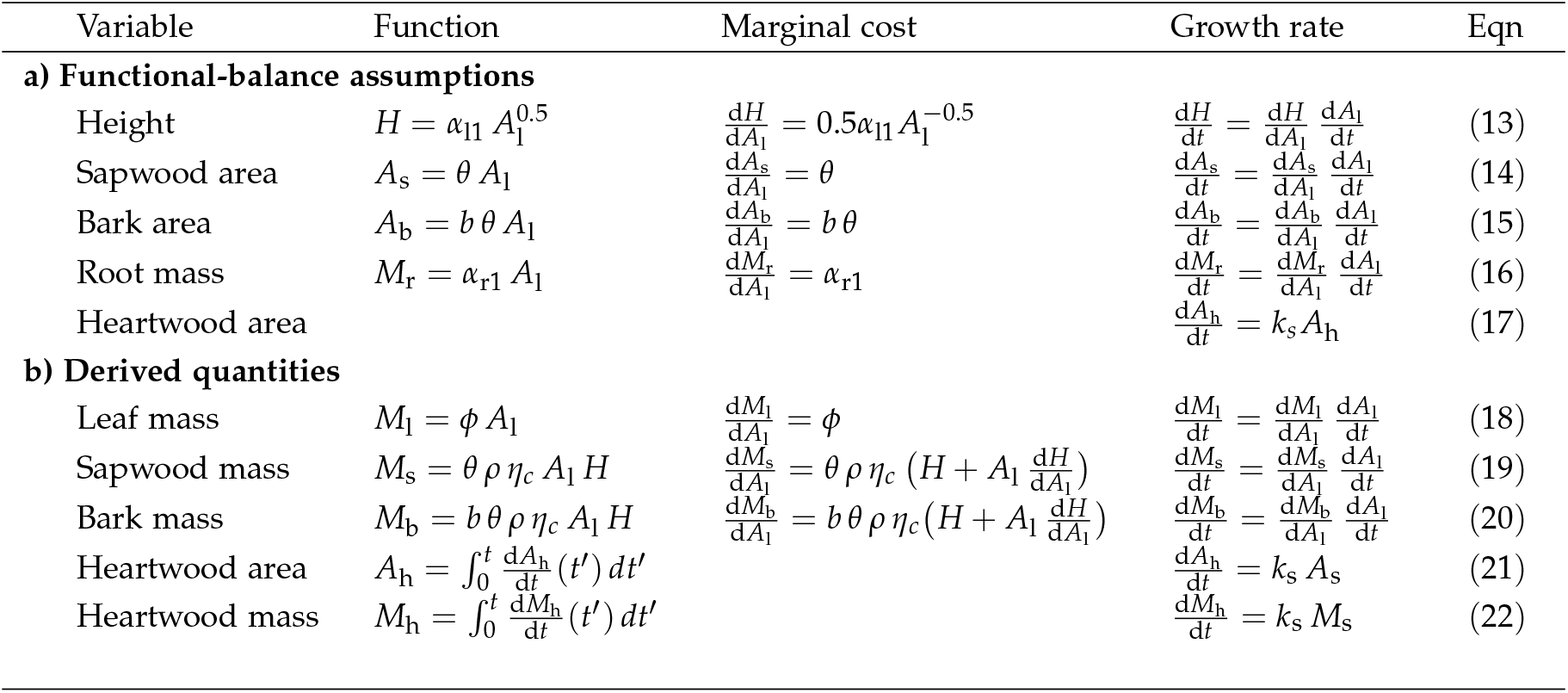
Equations for a “functional-balance” model of plant construction. The first column of part **a** provides cores assumptions between various size metrics and leaf area. Eqns in the middle and right columns of **b**, and in **b** can then be derived from the assumptions in the left column of **a**. The column “Eqn” indicates equation numbers referred to in the main text. See Table 2 for a list of variable names and definitions.

Assumption 1 accounts for the architectural layout of the plant. Assumptions 2-4 are realisations of the pipe model (Shinozaki *et al.*, 1964), whereby the cross-sectional area of conducting tissues are proportional to leaf area. To describe the vertical distribution of leaf area within the canopy of an individual plant (assumption 5), we use the model of Yokozawa & Hara (1995), which can account for a variety of canopy profiles through a single parameter *η*_*c*_, varying from 0-1 (for details, see Supplementary Materials).

Combined, the functional-balance assumptions from Table 4a lead directly to equations describing the mass of sapwood and bark in relation to leaf, and the amount of leaf in relation to height (Table 4b). Substituting from Table 4 into eqns 7, 8, 9 then gives all the necessary terms needed to implement the growth model described in Fig. 1.

### Trait-based trade-offs

We now consider how trait-based trade-offs enter into the above growth model. It is essential that any trait includes both a benefit and cost in terms of plant function and/or life history; otherwise we would expect ever-increasing trait values towards more beneficial values. For present purposes, we consider five prominent traits for which we can posit specific costs and benefits, outlined in Table 3. In postulating potential benefits and costs, we consider only those thought to arise as direct biophysical consequences of varying a trait. The trade-offs are then implemented as follows.

**Seedmass,** *ω*: There is a direct energetic trade-off between a plant’s fecundity and the size of its seeds. Moreover, we assume larger seeds result in larger seedlings; this seed-size number tradeoff translates into a demographic tradeoff between the number of seedlings produced by a parent and their initial size.
**Heightat maturation,** *H*_mat_: This trait moderates an inevitable energetic trade-off between growth and reproduction, that operates at all times through the lifestyle. Mass invested in growth cannot be invested in reproduction. To describe how the fraction of mass allocated to reproduction, 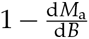, changes through ontogeny, we assume a function 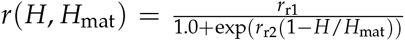, where *H*_mat_ is height at maturation, *r*_r1_ is the maximum possible allocation (0-1) and *r*_r2_ determines the sharpness of the transition. The exact shape of this function is non-critical, what is important is that plants shift from a period of investing mainly in growth to investing mainly in reproduction. The trait *H*_mat_ then describes the size at which this shift occurs, with direct biophysical consequences for growth.
**Nitrogenper unit leaf area,** *ν*: We allow for the maximum photosynthetic capacity of the leaf to vary with leaf nitrogen per unit area, as 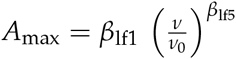, where *β*_lf1_, *ν*_0_ and *β*_lf5_ are constants. Respiration rates per unit leaf area are also assumed to vary linearly with leaf nitrogen per unit area, as *β*_lf4_ *ν*.
**Leafmass per unit area,** *φ:* This trait directly influences growth by changing 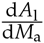 (Table 4). In addition, we link *φ* to the rate of leaf turnover (*k*_l_), based on a widely observed scaling relationship from Wright *et al.* (2004): 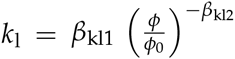 where *β*_kl1_, *φ*_0_ and *β*_kl2_ are empirical constants. Following Wright *et al.* (2004), the rate of leaf respiration per unit area is assumed independent of *φ*, as such the mass-based rate is adjusted accordingly whenever *φ* is varied.
**Wooddensity,** *ρ*: This trait directly influences growth by changing 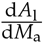 (Table 4). In addition, we link *ρ* to the rate of sapwood and bark turnover, mirroring the relationship assumed for leaf turnover: 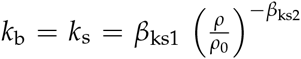 where *β*_ks1_, *ρ*_0_ and *ρ*_ks2_ are empirical constants. The rate of sapwood and bark respiration per unit stem volume is assumed to be independent of *ρ*, as such the mass-based rate is adjusted accordingly whenever *ρ* is varied. There is very little data on rates of sapwood turnover and respiration in relation to wood density, so this latter assumption is more speculative than the equivalent assumption for leaves, which is well supported empirically.

## Methods

The growth model described above has been implemented as the `FF16` physiological model within the `plant` package (Falster *et al.*, 2016) for R (R Core Team, 2015). The `plant` package also makes use of supporting packages `Rcpp` (Eddelbuettel, 2013) and the Boost Library for C++(Schäling, 2014), via the package `BH` (Eddelbuettel *et al.*, 2015). To encode the trait-based trade-offs described above, we use the capacity to in `plant` to provide a “hyper-parameterisation” function, which enables various parameters to covary with traits (for full details see Supplementary Materials – A).

For the most part, parameters used in the current analysis were sourced from Falster *et al.* (2016) (see Tables S1-S2 for values). The only exceptions are: i) parameters affecting the relationships outlined in Table 4a, which were estimated from data described below, and ii) parameters describing the function for reproductive allocation. By default, we set *r*_r1_ = 0.8 and *r*_r2_ = 10, implying a relatively rapid transition to reproduction at *H* = *H*_mat_ (see panel for 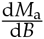 in Fig. 1).

The functional-balance assumptions listed in Table 4a were evaluated using data from the Biomass and Allometry Database (BAAD) (Falster *et al.*, 2015), which includes records for various size metrics from 21084 individual plants across 656 species. We fit standardised major axis (SMA) lines (Warton *et al.*, 2006) to characterise bivariate relationships. We implemented a hierarchical model structure, where the distribution of slopes and intercepts among groups is assumed to come from normal distributions. The means and variances of these distributions are then fit as part of the model-fitting procedure.

The analyses presented employ best practises in scientific computing, as defined by Wilson *et al.* (2014), and are fully reproducible, via code available at github.com/traitecoevo/growth_trajectories.

## Results

### Model assumptions

To verify model assumptions we compared the assumptions outlined in Table 4a to data sourced from the BAAD (Falster *et al.*, 2015). Additionally, we evaluated an important prediction arising from the eqns in Table 4a, that the amount of live stem tissue supporting each unit of leaf area increases linearly with height, as per the equation

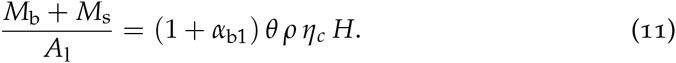

Note that *α*_b1_, *θ*, *ρ*, *η*_*c*_ are all traits, i.e. properties that are assumed approximately constant through ontogeny for any given species. Fig. 3 shows that the three functional-balance assumptions outlined in Table 4a and the relationship in eqn 11 are all well-supported by the available data. The solid lines indicate SMA lines fit to each species in the dataset. Dashed lines indicate the slope of predictions under functional-balance assumptions in Table 4a and eqn 11. As expected, species differed in elevation, but less so in the slope of the fitted lines; with slopes aligning with those predicted by the functional-balance assumptions.

**Figure 3:**
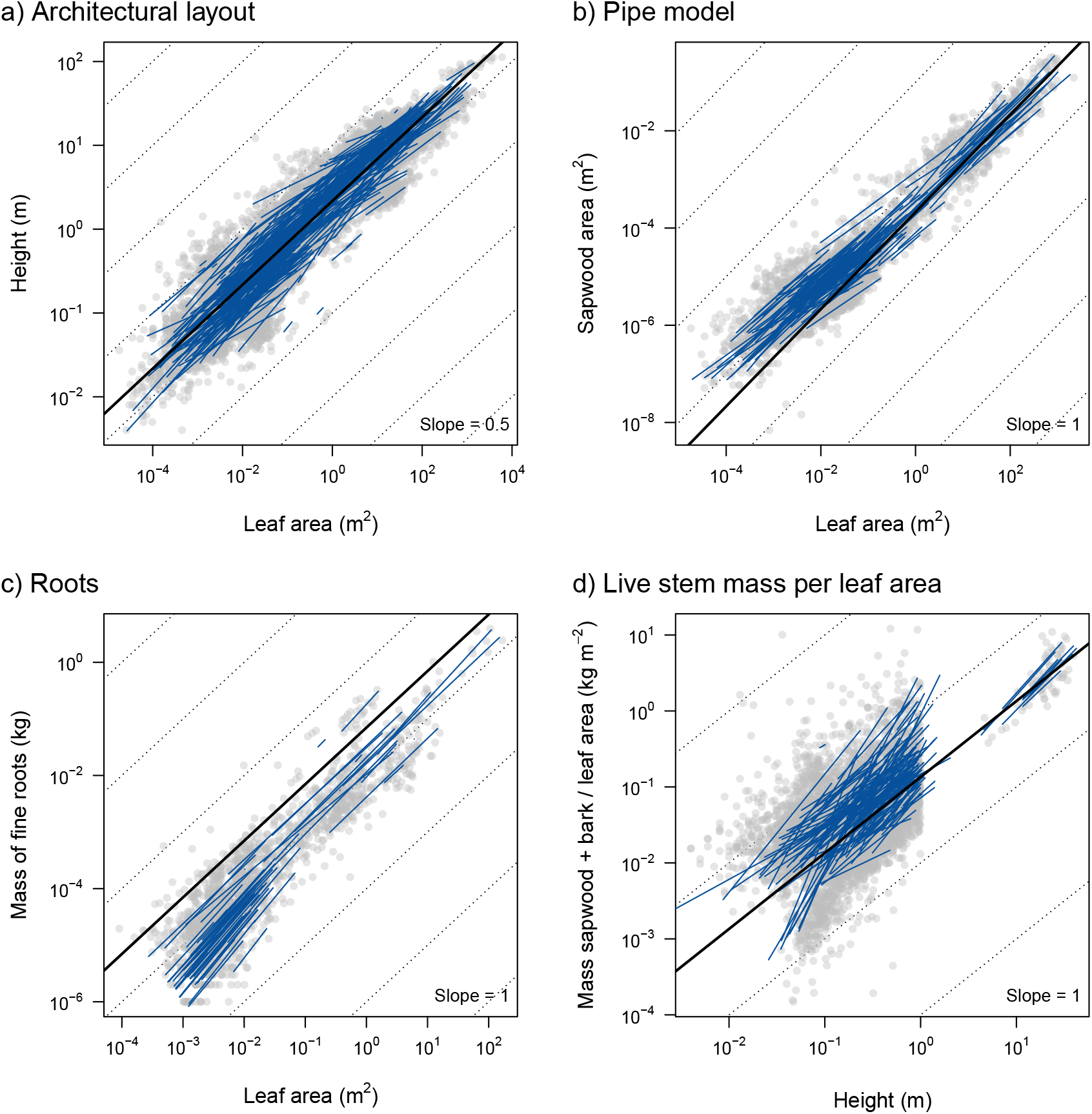
Key assumptions of a functional balance model for plant construction. Each dot is a single plant from the BAAD (Falster *et al.*, 2015). Blue lines indicate the show standardised major axis of the bivariate cloud for each species. Dotted lines indicate slope of predictions under functional-balance assumptions in Table 4. The dark line shows the relationship assumed in the plant model and applied throughout the paper.

### Changes in growth rate with size

Our growth model suggests an intrinsically size-dependent pattern of biomass-production and growth, which aligns with well-known empirical patterns (Table 1). The panels in Fig. 1 show the expected patterns for a typical woody plant, obtained by applying the functional-balance model encoded in `plant`. Biomass production shows a hump-shaped pattern with size, decreasing at larger sizes as the turnover and respiration of sapwood and bark increase. Height growth also shows a hump-shaped pattern with size, first increasing then decreasing. This pattern results from systematic changes in the four components of eqn 7 with size, including a strong decline in the fraction of plant that is leaf declines with increasing size (Fig. 1), increasing reproductive allocation (Fig. 1), and declining mass production. In contrast, basal-area growth continues to increase with size, due to an increasing influence of stem turnover. Diameter growth shows a weakly hump shaped curve, tapering off slightly at larger sizes, in part because of the allometric conversion from basal area to diameter (eqn 9, and in part because of increased reproductive allocation in older trees (Fig 1). All growth measures decrease sharply with size when expressed as relative growth rates (results not shown).

### Changes in height growth rate with traits

We analysed the response of growth rate to five different traits under the assumed trade-offs (Table 3). We considered changes in absolute and relative growth rates for mass, height, stem area and stem diameter. The first two traits considered modify behaviour primarily at the start and end of an individual’s growth trajectory, and are therefore termed “ontogenetic traits”. The remaining three traits are termed “development traits” because they moderate the rate of movement along an individual’s growth trajectory. Across the five different traits, we observed four relatively distinct types of response. These responses are summarised in Table 5 and described in more detail below.

**Table 5:** Predicted effects of traits on key elements of plant function determining growth rate. Arrows indicate the effect an increase in trait value would have on each element of the equations, with dashes indicating no effect. Traits are: seed mass (*ω*), height at maturation (*H*_mat_), nitrogen content per leaf area (*ν*), leaf mass per unit leaf area (*φ*), and wood density (*ρ*) (see Table 3 for more details). Adapted and expanded from Gibert et al. (2016).

|  | Symbol | Ontogenetic |  | Development |  |  |
| --- | --- | --- | --- | --- | --- | --- |
| | | $\omega$ | $H_{\text{mat}}$ | $\nu$ | $\phi$ | $\rho$ |
| a) Effect on elements of eqns 1 – 9 |  |  |  |  |  |  |
| Net biomass production | $\text{dB}/\text{dt}$ | | | | | |
| • Photosynthesis |  | - | - | ↑ | - | - |
| • Respiration |  | - | - | ↑ | - | - |
| • Turnover |  | - | - | - | ↑ | ↑ |
| Allocation to growth | $\text{dM}_a/\text{dB}$ | - | ↑ | - | - | - |
| Leaf deployment per mass | $\text{dA}_l/\text{dM}_a$ | | | | | |
| • Leaf |  | - | - | - | ↓ | - |
| • Sapwood |  | - | - | - | - | ↓ |
| • Root |  | - | - | - | - | - |
| Architecture | $\text{dH}/\text{dA}_l$ | - | - | - | - | - |
| b) Predicted effect of trait on growths rate for a small and large plant |  |  |  |  |  |  |
| Height |  |  |  |  |  |  |
| • Absolute | $\text{dH}/\text{dt}$ | ↑- | -↑ | ↑↑ | ↓- | ↓↓ |
| • Relative | $\text{dH}/(\text{dt.H})$ | ↓- | -↑ | ↑↑ | ↓- | ↓↓ |
| Stem area |  |  |  |  |  |  |
| • Absolute | $\text{dA}_{\text{st}}/\text{dt}$ | ↑- | -↑ | ↑↑ | ↓- | ↓↓ |
| • Relative | $\text{dA}_{\text{st}}/(\text{dt.A}_{\text{st}})$ | ↓- | -↑ | ↑↑ | ↓- | ↓↓ |
| Stem diameter |  |  |  |  |  |  |
| • Absolute | $\text{dD}/\text{dt}$ | ↑- | -↑ | ↑↑ | ↓- | ↓↓ |
| • Relative | $\text{dD}/(\text{dt.D})$ | ↓- | -↑ | ↑↑ | ↓- | ↓↓ |
| Mass |  |  |  |  |  |  |
| • Absolute | $\text{dM}_t/\text{dt}$ | ↑- | -↑ | ↑↑ | ↓- | ↓↓ |
| • Relative | $\text{dM}_t/(\text{dt.M}_t)$ | ↓- | -↑ | ↑↑ | ↓- | ↓↓ |

#### Ontogenetic traits

**Seed size,** *ω*. Increasing *ω* in our model causes seedlings to be larger and fecundity to decrease. As such, the only effect of seed size on growth comes from changing the plant’s initial size. The plots in Fig. 1, which show changes in growth rate with plant size, also express the expected changes in the growth of seedlings due to changes in seed size. Under similar light conditions, larger seedlings are predicted to have faster absolute growth rates in all metrics because of their greater total leaf area. At the same time, relative growth rate is predicted to decrease with size, because the ratio of leaf area to support mass decreases with plant size. As plants grow, differences in initial mass will decrease in importance, relative to other factors influencing growth through the life-cycle. As a result, the correlations between seed size and growth rate observed for seedlings disappear among larger plants.

**Height at maturation,** *H*_mat_. *H*_mat_ moderates growth by adjusting the amount of energy invested in growth, i.e. the term 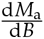 in eqns 7 and 8. Greater *H*_mat_ can thus lead to a growth advantage by increasing 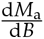 (Fig. 4). At smaller sizes, there is no differentiation among species, because most individuals are focusing on growth. At larger sizes, individuals of some species are allocating a larger fraction of biomass to reproduction, which reduces their growth rate relative to those species with greater *H*_mat_ (Fig. 4).

**Figure 4:**
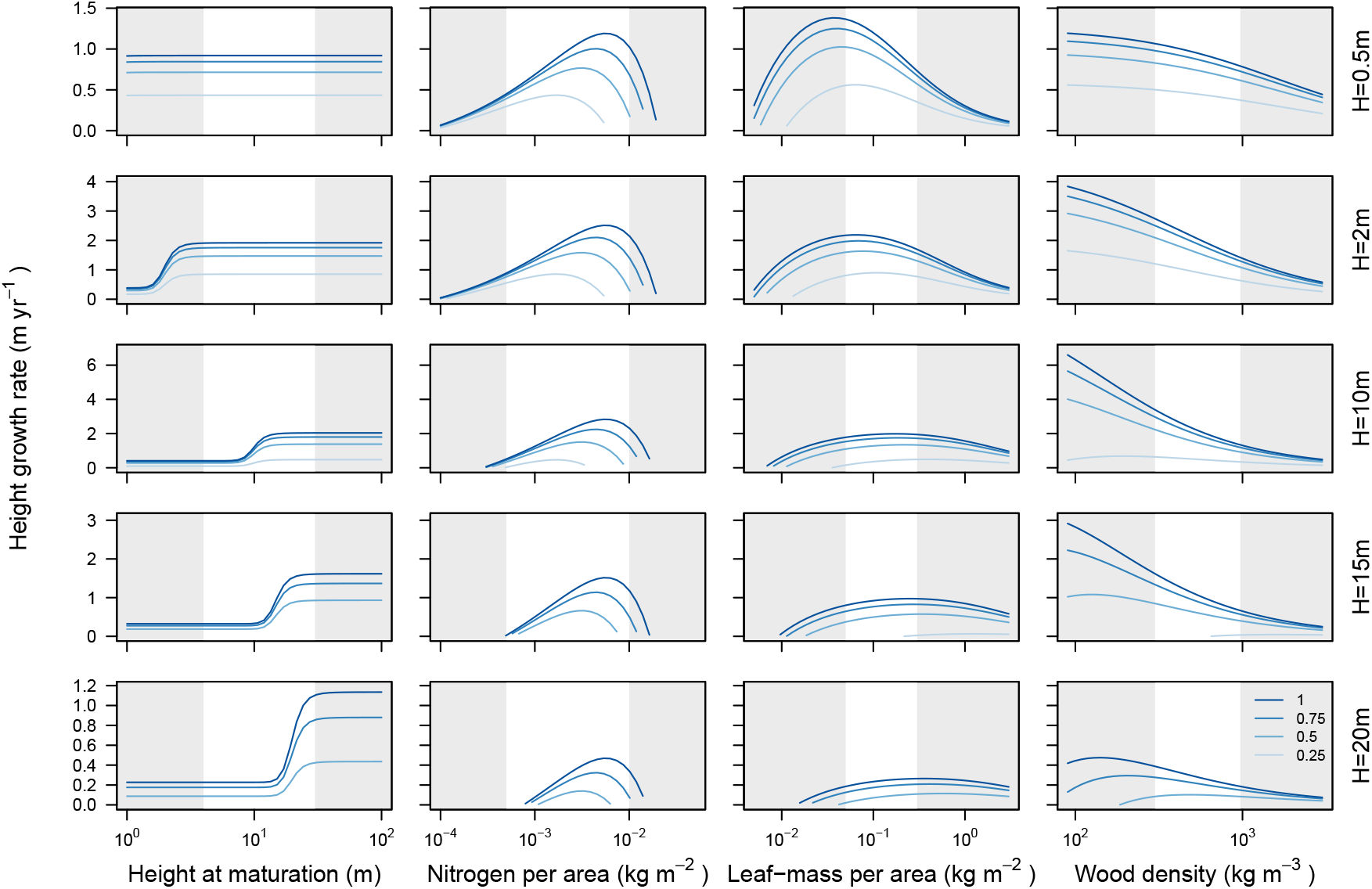
Effect of four traits on height growth rate for different-sized plants. Growth rates were simulated using the `plant` model, applying the trade-offs describing in Table 3. Each panel shows how growth is influenced by a different trait for plants of a given height, and across a series of canopy openness values from completely open (light blue, *E* = 1) to heavily shaded (dark line, *E* = 0.25). For any given value of trait and *E*, plants were grown to the desired height and their growth rate estimated. The white regions indicate trait ranges that are typically observed in real systems. Figs. S2-S4 show similar plots but with growth measured as stem diameter, stem area, or plant mass. Changes in trait-growth relationships are summarised in Table 5.

#### Development traits

The remaining three traits (*ν*, *φ*, & *ρ*) moderate growth rates at a point along an individual’s growth trajectory (Fig. 4).

**Leaf nitrogen content per leaf area,** *ν*. The response of growth rate to changes in *ν* is relatively straightforward: there is an optimum value of *ν* that maximises height growth rate in a given light environment *E* and does not vary with height (Fig. 4). As *E* increases from low to high, the optimal *ν* also increases. The invariance of the growth-trait relationship with respect to size arises as follows. The direct physiological effect of *ν* is to increase the maximum potential photosynthetic rate of leaves, with a cost of higher respiration rate. Both the cost and benefits of *v* appear within 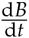, implying the direction of correlation between trait and growth rate depends crucially on the change in mass production per *ν*. From eqn 1, 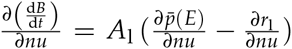. Since both and 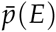 and *r*_l_ are expressed per unit area and independent of height, the optimal value is also independent of plant height.

**Leaf mass per unit leaf area,** *φ*. Unlike *ν*, the response of growth rate to changes in *φ* varies strongly with plant height, with the relationship moving like a wave across the trait spectrum (Fig. 4). As a result, the value of *φ* that optimises plant growth increases with plant size, and the direction of correlation between height growth rate and *φ* shifts from negative to positive, as plants increase in height. Decreasing *φ* has two impacts on height growth rate. First, lower *φ* increases the leaf deployment per mass invested 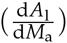 by economising on construction costs. Second, lower *φ* decreases net production 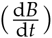, due to increased leaf turnover. Whether lower *φ* increases growth thus depends on the relative magnitude of these two effects. When plants are small the effect on leaf deployment rate is larger and so decreasing *φ* increases growth rate. When plants are large, the influence of *φ* on leaf deployment rate is diminished, because the cost of building other supportive tissues (other terms in eqn 5) is larger (Fig. 1). The net result is that at larger sizes, low *φ* is no longer advantageous for growth (Fig. 4).

**Wood density,** *ρ*. As for leaves, lower cost of stem construction (lower *ρ*) decreases the cost of deploying a unit of leaf area, and may thereby increase growth rates (Fig. 4). In contrast to *φ*, the benefits of cheaper stem construction become more pronounced at larger sizes, as an increasingly large fraction of the plant is wood (Fig. 1a). Thus, *ρ* has only a weak effect on growth rates for small plants, but a strong effect for intermediate to large plants. As the plant becomes very tall (H >15m), the benefits of low *ρ* finally begin to diminish.

### Changes in other growth rates with trait

The results reported above and shown in Fig. 4 focus on height growth rate (eqn 7). Corresponding results for growth rates in stem diameter (eqn 9), stem basal area (eqn 8), and above-ground mass (eqn 4) are provided in Figs. S2-Fig. S4 of the Supplementary Materials. In each figure, plants were grown to a suitable diameter, area, or mass. As such, changes in relative growth rates with traits show a similar patterns as absolute growth rates.

We find that for seed mass, leaf nitrogen, leaf mass per area and height at maturation, the patterns of growth rate in stem diameter, stem area, or above-ground mass with respect to traits mirror those observed with respect to heigh growth (Table 5). The only trait where a slightly different response was observed was for wood density. Whereas the effect of wood density on height growth tended to diminish slightly at larger sizes (Fig. 4), the effect became even stronger when measuring growth rate in stem diameter, stem area or above-ground mass. Recall that sapwood lost via turnover is turns into heartwood. Whereas the loss of sapwood diverts energy away from height growth rate, the faster accumulation of heartwood actually accelerates the growth of stem diameter and area.

### Responsiveness of growth rate to light

The predictions in Figs. 4 and S2-S4 illustrate how traits impact on growth rate under different light environments and at different sizes. An additional outcome that arises directly from these analyses is that traits moderate the responsiveness of growth to changes in light environment. This response arises because individuals with higher potential growth rate naturally have greater potential plasticity in growth. Our results therefore support findings that species with low *ρ* increase growth more substantially with increases in light (Table 1). Variation in *φ* also moderates the response of growth to changes in light, with species having the lowest *φ* being most responsive. However, unlike for *ρ*, the effect appears only for the smallest size classes.

### Shade tolerance

Combining eqn 10 with the function-balance model in Table 4 leads to the a more specific expression for calculating WPLCP, as the value of *E*\* that gives

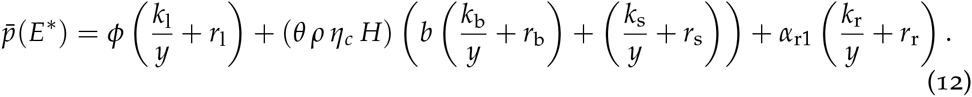

Eqn 12 indicates WPLCP will increase approximately linearly with *H* and potentially vary with traits *ν*, *φ*, and *ρ*. With some further manipulations, it is possible to show that WPLCP will decrease with *φ* if *β*_kl2_ > 1. Likewise WPLCP will decrease with *ρ* if *β*_ks2_ > 1. The parameters *β*_kl2_ and *β*_ks2_ give the slope relating tissue turnover rate to *φ* and *ρ*, respectively. Since in this analysis, we have assumed these criteria to hold, species with low *φ* and low *ρ* are predicted to be less shade tolerant because of disproportionate increases in turnover costs (Fig. 5). At low *φ*(*ρ*), leaf (sapwood) turnover is higher and thus a greater light income is needed to offset these costs. WPLCP also decreases with height because as size increases, the total amount of carbon needed to offset respiratory and turnover costs in the stem also increases (Givnish, 1988). In addition, WPLCP varies with *ν*. At small sizes, WPLCP increases with *ν* across the band of values typically observed in real plants, i.e. high leaf nitrogen makes seedlings shade intolerant. At larger sizes, as net production declines to zero, WPLCP begins to increase again for very low values of *ν*. All of these patterns match empirically observed patterns (Table 1).

**Figure 5:**
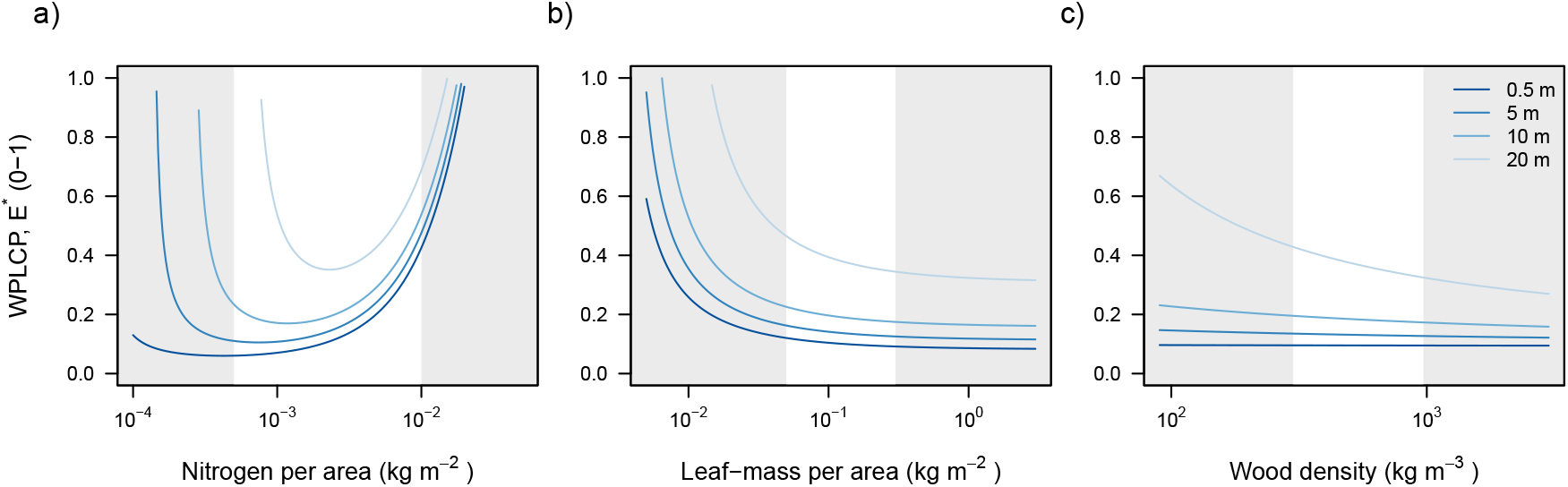
Effect of three development traits on shade tolerance. Panels show effect of traits on level of canopy openness that causes net production (eqn 1) to be zero. Different lines indicate relationship for plants with specified height, from short (light blue, *H* = 0.5m) to tall (dark line, *H* = 20m). The white regions indicate trait ranges that are typically observed in real systems.

## Discussion

Using a model relating plant physiological function and carbon allocation to five prominent traits, we have shown how traits impact on plant growth across the life cycle. This approach extends a widely-used theoretical model for seedlings, which links mass-based growth rate to the trait leaf mass per unit leaf area (Lambers & Poorter, 1992; Wright & Westoby, 2000), to explicitly include influences of size, light environment, and other prominent traits. During the last two decades, functional traits have captured the attention of ecologists, in large part because of the ability to organise the world’s plant species along standard dimensions (Westoby *et al.*, 2002). However, it has remained unclear how or whether prominent traits influence growth outcomes (Poorter *et al.*, 2008; Wright *et al.*, 2010; Paine *et al.*, 2015). Matching a growing amount of empirical evidence (Table 1), this study outlines when and why the direction or strength of correlation between traits and growth rate shifts with plant size. Moreover, we show that different traits and trade-offs generate different types of response. Combined with the available empirical evidence, these results demand a fundamental shift in our understanding of plant ecological strategies, away from one in which species are thought to have a fixed growth strategy throughout their life (from slow to fast growth) (e.g. Grime, 1977; Adler *et al.*, 2014; Paine *et al.*, 2015) to one in which traits define a size-dependent growth trajectory (Gibert *et al.*, 2016). Moreover, we find that growth trajectories and the ranking of traits across them are also moderated by the light environment; while traits that minimise costs of tissue respiration and / or turnover also make plants more shade tolerant (i.e. lower WPLCP), as is empirically observed (Messier *et al.*, 1999; Craine & Reich, 2005; Poorter & Bongers, 2006; Baltzer & Thomas, 2007; Lusk *et al.*, 2008). The plant model, used here, builds on and extends several related approaches, wherein emergent outcomes such as height, diameter and mass growth arise from the interaction of different tissues and traits (e.g. Givnish, 1988; Mäkelä, 1997; Moorcroft *et al.*, 2001). This approach is quite different to models derived from metabolic scaling theory (MST), which derive everything from a single master “scaling” equation for mass growth and have thus far been unable to account for size-dependent changes in the correlation between traits and growth rate (Enquist *et al.*, 1999, 2007). Our approach is also fundamentally different from statistically-fitted growth models (e.g. Hérault *et al.*, 2011; Rüger *et al.*, 2012; Iida *et al.*, 2014) in that it predicts rather than statistically tests for trait-based effects. In this sense, our model is designed to both explained observed phenomena (Table 1) and also generate new hypotheses.

### Generalising to other traits and trade-offs

The model presented here extends a widely-used theoretical model for seedlings, which links mass-based growth rate to the trait *φ* (Lambers & Poorter, 1992; Wright & Westoby, 2000), to larger plants and other traits. Importantly, the seedling model can be derived as a special case of our extended approach. Unlike the original model for seedlings (eqn 2), the new model also predicts a relationship between *φ* and growth rate that changes with size. In particular, the correlation shifts from being strongly negative in seedlings to being absent, or even possibly positive in larger plants (Fig. 4), irrespective of whether growth rate is estimate via height, stem diameter, stem area or total mass. This shift, which matches empirical evidence (Poorter *et al.*, 2008; Wright *et al.*, 2010; Hérault *et al.*, 2011; Paine *et al.*, 2015; Gibert *et al.*, 2016), occurs because the benefits of cheap leaf deployment diminish with plant size. As seedlings, leaves comprise a large part of the plant (Fig. 1a). Decreasing *φ* then has an overwhelmingly positive effect on height and diameter growth rate because the effect of increasing 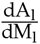 is large compared to the other terms in eqn 5. As plants increase in size, however, the amount of supporting tissue increases (Fig 3d), decreasing the benefit of cheaper leaf construction in eqn 5. Consequently, the effect of *φ* on leaf turnover comes to dominate at larger sizes, and as such, the effect of *φ* on height, diameter and, mass growth shifts from negative to either flat or mildly positive (Table 5).

The list of functional traits that are known to differ among plant species is long and ever-increasing (Pérez-Harguindeguy *et al.*, 2013). While we have focussed on understanding the effects of five specific traits on growth rate, the framework presented can be extended to generate hypotheses about other traits and tradeoffs. The main criteria for including new traits is that a clear trade-off has been established, with benefits and/or costs that ultimately translate into biomass or carbon, and can therefore be embedded within the eqns in Fig. 1b. While the list of plant traits that have been measured is extensive, clear trade-offs have been established for only a few of these. A well-developed trade-off must include two opposing forces, that operate at some point in the organisms life cycle. It is not necessary that benefit and cost both enter into Fig. 1; for example wood density is sometimes viewed as a trade-off between the costs of tissue construction and the rate of stem mortality. In that case, the costs of lower wood density would not appear within the eqns in Fig. 1b, so lower wood density would always increase growth rate.

Our framework also highlights what is needed for traits to impact on growth rate and shade tolerance. While traits can influence many aspects of plant function, these influences must operate via the pathways outlined in Fig. 1 if the trait is going to impact on growth. For example, many studies have focused on traits related to plant hydraulics, such as vessel size and increased sapwood area per leaf area (Zanne *et al.*, 2010). These traits will inevitably influence the rate of photosynthesis per leaf area (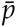 in eqn 1) by altering conductance of water to the leaf. The potential costs of larger vessel size might be higher rates of stem turnover, which would appear in the term *k*_s_ in eqn 1. The costs of increased sapwood area per leaf area is increased allocation to stem, a factor which is already included in our framework via the parameter *θ* (Table 4). The effect of both these traits on growth rate should be expected to vary with plant size.

### Implications for trait-based approaches

There are some broad implications of our work for our understanding of plant ecological strategies and plant growth.

First, our results highlight the importance of allocation decisions and turnover costs in determining growth dynamics. Much of current ecosystem research focusses on factors effecting primary production – photosynthesis, respiration, and resultant fluxes of carbon – with less attention devoted to allocation and turnover (Friend *et al.*, 2014; for comparisons of models see Sitch *et al.*, 2008; De Kauwe *et al.*, 2014). Yet, for four of the five traits considered here, trait values do not influence net primary production. In fact, the analysis with *φ* shows that increased growth rate can occur even at a distinct cost to the plant’s carbon budget. Low *φ* results in high leaf turnover, such that individuals with a *φ* have lower mass production. It is this property that makes them shade intolerant. And yet they can still achieve a growth advantage (when small), because the benefits of cheap leaf construction outweigh the costs of high leaf turnover.

Second, our results demand a shift in the way plant species are sometimes described as, being of fast or slow growth. To the extent the ranking of growth rates among individuals differing in traits shifts with either plant size or light environment, it is not possible to describe a species via a single point along a spectrum from slow to fast growth. Such a spectrum is implied by many of theoretical models used in community ecology, including Grime’s CSR triangle, the r-K spectrum, and coexistence models base on the Lotka-Volterra system of equations (e.g. Grime, 1977; Chesson, 2000). Researchers using functional traits have also tended to describe species as fast or slow growing (e.g. Adler *et al.*, 2014; Díaz *et al.*, 2016). Our results suggest a more-nuanced approach. Plants that are fast growing as seedlings may not be fast growing as saplings or adults, or under low light. Plants that are fast growing as adults may not be fast growing as seedlings. This more-nuanced perspective tends to mirror observed demographic patterns, where juvenile and adult growth rates are sometimes only loosely correlated (Rees *et al.*, 2001).

Third, our results suggest that even if traits define a potential growth trajectory, researchers seeking to link traits to growth rate must probe deeper into the data than simply looking for a linear relationship between traits and average growth, to recover the expected relationships. None of the predicted relationships between traits and growth is linear across the range of sizes and light environments tested. As such, we should not be surprised if the mean growth rate across individuals spanning a range of sizes or light environments is only weakly or not correlated with traits (e.g. Poorter *et al.*, 2008; Paine *et al.*, 2015). Controlling for size, site and light environment will be essential for detecting significant patterns (e.g. Gibert *et al.*, 2016), as will having a clear expectation for the hypothesised relationship.

While our theory has succeeded in explaining some observed phenomena (Table 1), the test for good theory is that it also makes new predictions that enable the theory to be further refined and tested. To that end, we can make a further prediction arising from our results, which is that the trait *φ* should increase through ontogeny for all individuals, across all species. Such shifts have been observed across a variety of species King (1999); Thomas & Bazzaz (1999); Koch *et al.* (2004). Since the value of cheap leaf construction diminishes with size, it pays for plants – and especially those with low *φ* – to increase their *φ* as they grow larger. King (1999) made a similar prediction for a single species of *Eucalyptus*, but here we can extend the idea across species. While trait-based research largely focusses on differences among species, it has long been recognised that traits also vary among individuals within a species and within individuals (Westoby *et al.*, 2002). This hypothesis attempts to give meaning to some of that variation, and shows how variation across and within species might be understood within a single framework.

### Comparison with other frameworks

As noted above, the `plant` model is closely related to models used in several other studies, including those by Givnish (1988); Yokozawa & Hara (1995); Mäkelä (1997); King (1999, 2005); Moorcroft *et al.* (2001); Li *et al.* (2014), and in particular those by Mäkelä (1997) and Moorcroft *et al.* (2001). These models have several properties in common, including that they all have growth being driven by the gross amount of photosynthetic income; have photosynthesis increasing non-linearly with light and leaf nitrogen content; and that they consider the costs of respiration and turnover in different tissues. Many models also make functional-balance assumptions, for example linking the cross-section of sapwood to leaf area (Givnish, 1988; Yokozawa & Hara, 1995; Mäkelä, 1997; King, 2005; Moorcroft *et al.*, 2001). We note that an assumption of exact functional balance is not critical for our results, what matters is that the amount of live biomass (i.e. excluding heartwood) needed to support an extra unit of leaf area increases with height (as Fig. 3d). Some models also differ from in that they directly link a plant’s stem diameter to its height, e.g. Yokozawa & Hara (1995); King (1999); Moorcroft *et al.* (2001); Li *et al.* (2014), whereas we let this scaling arise as an emergent outcome of growth and sapwood turnover. Again, we expect this difference will not affect the main results.

A feature distinguishing our model from most of those mentioned above is the explicit linking to trait-based trade-offs. While such a linkage was also made by Moorcroft *et al.* (2001) in the ED model; analyses using ED have mainly focussed on ecosystem-level outcomes rather than the growth of individual plants. Because of its underlying similarities, we expect the dynamics reported here to be also present within the ED model. King (1999) also connected his model of growth for a single species to the trait *φ*, and like the current study, predicted a gradual flattening out of the relationship between *φ* and growth rate with size (as in Fig. 4), because the influence of cheap leaf construction decreased with size. In our study however, there was additional cost of increased leaf turnover, that further penalised low *φ* strategies when plants were large.

Perhaps the two most controversial elements of our approach concerns the assumptions about tissue replacement and reproductive allocation. Many vegetation models determine allocation based on net primary production (photosynthesis - respiration), whereas we also subtracted tissues lost via turnover before distributing surplus biomass. This is because we assume tissues lost via turnover are replaced before carbon is allocated to either new growth (i.e. growth that leads to a net increase in *M*_l_, *M*_b_, *M*_s_ or *M*_r_) or reproduction (Thornley & Cannell, 2000). This assumption is likely to hold true for most woody plants and perennials, but may not hold for some herbs or annuals, where the switch to reproduction may entail a run-down in the vegetative part of the plant. The second assumption we make is that when mature, plants allocate a substantial fraction of their surplus carbon to reproduction. While it remains unclear just how much adult plants might allocate to reproduction, recent reviews suggest the fraction may be high (Thomas, 2011; Wenk & Falster, 2015). Moreover, a long line of theoretical models indicate that allocation should increase as plants age (reviewed by Wenk & Falster, 2015). Reproductive allocation is given little attention in models of ecosystem flux (e.g. Sitch *et al.*, 2008; De Kauwe *et al.*, 2014). For example in the ED model, a fixed 30% of net primary production is allocated to reproduction, irrespective of plant size. In the case of growth, differences in reproductive allocation offer a clear pathway for explaining patterns linking a plants maximum size to the growth rate of large individuals (e.g. Wright *et al.*, 2010).

Another class of model dealing explicitly with size-related effects includes those derived from the metabolic scaling theory (MST) of ecology (Enquist *et al.*, 1999, 2007). Several points suggest our new framework provides a better explanation to the growth phenomena in Table 1 than the MST framework. First, the MST-derived model suggests diameter growth continues to increase as plants grow, whereas empirical data suggests growth rate declines for larger plants (Canham *et al.*, 2004, 2006; Hérault *et al.*, 2011). Second, the MST model does not allow for the effects of traits to vary with plant size. Predicted effects are for a linear increase in growth with decreases in either *φ* and lower *ρ*, that apply irrespective of size. However, at least for *φ* such effects in large trees have not been observed.

### Closing remarks

Overall we have shown how diverse phenomena related to plant growth can be understood with a model accounting for processes generating photosynthetic in-come and allocating this among different tissues. The need to consider effects of plant size, alongside trait-based differences among species, has has long been recognised in trait-based research (e.g. Givnish, 1988; Thomas & Bazzaz, 1999; Moorcroft et al., 2001; Westoby et al., 2002; Enquist et al., 2007). Here we have provided a framework for achieving this. By disentangling the effects of plant size, light environment and traits on growth rates, our results provide a solid theoretical foundation for trait ecology and thus provide a platform for understanding growth across diverse species around the world.

## Acknowledgements

We thank J Camac, A Gibert, G Kunstler, C Prentice, E Wenk, M Westoby, I Wright, and SJ Wright for helpful discussions; J Camac for introducing us to the `stan` framework, and D Warton for discussions about SMA line fitting. Falster was supported by an Australian Research Council discovery grant (DP110102086). FitzJohn was supported by the Science and Industry Endowment Fund (RP04-174). The authors have no conflicts of interest to declare.

## Supplementary Materials

A – Additional details on the `FF16` growth model
B – Supplementary Tables
C – Supplementary Figures

## References

1. Adler, P.B., Salguero-Gomez, R., Compagnoni, A., Hsu, J.S., Ray-Mukherjee, J., Mbeau-Ache, C. & Franco, M. (2014). Functional traits explain variation in plant life history strategies. Proceedings of the National Academy of Sciences, 111, 740–745.

2. Anderson-Teixeira, K.J., Davies, S.J., Bennett, A.C., Gonzalez-Akre, E.B., Muller-Landau, H.C., Wright, S.J., Salim, K.A., Zambrano, A.M.A., Alonso, A., Baltzer, J.L., Basset, Y., Bourg, N.A., Broadbent, E.N., Brockelman, W.Y., Bunyavejchewin, S., Burslem, D.F.R.P., Butt, N., Cao, M., Cardenas, D., Chuyong, G.B., Clay, K., Cordell, S., Dattaraja, H.S., Deng, X., Detto, M., Du, X., Duque, A., Erikson, D.L., Ewango, C.E., Fischer, G.A., Fletcher, C., Foster, R.B., Giardina, C.P., Gilbert, G.S., Gunatilleke, N., Gunatilleke, S., Hao, Z., Hargrove, W.W., Hart, T.B., Hau, B.C., He, F., Hoffman, F.M., Howe, R.W., Hubbell, S.P., Inman-Narahari, F.M., Jansen, P.A., Jiang, M., Johnson, D.J., Kanzaki, M., Kassim, A.R., Kenfack, D., Kibet, S., Kinnaird, M.F., Korte, L., Kral, K., Kumar, J., Larson, A.J., Li, Y., Li, X., Liu, S., Lum, S.K., Lutz, J.A., Ma, K., Maddalena, D.M., Makana, J.R., Malhi, Y., Marthews, T., Serudin, R.M., McMahon, S.M., McShea, W.J., Memiaghe, H.R., Mi, X., Mizuno, T., Morecroft, M., Myers, J.A., Novotny, V., de Oliveira, A.A., Ong, P.S., Orwig, D.A., Ostertag, R., den Ouden, J., Parker, G.G., Phillips, R.P., Sack, L., Sainge, M.N., Sang, W., Sri-ngernyuang, K., Sukumar, R., Sun, I.F., Sungpalee, W., Suresh, H.S., Tan, S., Thomas, S.C., Thomas, D.W., Thompson, J., Turner, B.L., Uriarte, M., Valencia, R., Vallejo, M.I., Vicentini, A., Vrska, T., Wang, X., Wang, X., Weiblen, G., Wolf, A., Xu, H., Yap, S. & Zimmerman, J. (2015). CTFS-ForestGEO: a worldwide network monitoring forests in an era of global change. Global Change Biology, 21, 528–549.

3. Baltzer, J.L. & Thomas, S.C. (2007). Determinants of whole-plant light requirements in Bornean rain forest tree saplings. Journal of Ecology, 95, 1208–1221.

4. Blackman, V.H. (1919). The compound interest law and plant growth. Annals of Botany, os-33, 353–360.

5. Canham, C.D., LePage, P.T. & Coates, K.D. (2004). A neighborhood analysis of canopy tree competition: Effects of shading versus crowding. Canadian Journal of Forest Research, 34, 778–787.

6. Canham, C.D., Papaik, M.J., Uriarte, M., McWilliams, W.H., Jenkins, J.C. & Twery, M.J. (2006). Neighborhood Analyses Of Canopy Tree Competition Along Environmental Gradients In New England Forests. Ecological Applications, 16, 540–554.

7. Castro-Diez, P., Puyravaud, J.P., Cornelissen, J.H.C. & Villar-Salvador, P. (1998). Stem anatomy and relative growth rate in seedlings of a wide range of woody plant species and types. Oecologia, 116, 57–66.

8. Chave, J., Coomes, D.A., Jansen, S., Lewis, S.L., Swenson, N.G. & Zanne, A. (2009). Towards a worldwide wood economics spectrum. Ecology Letters, 12, 351–366.

9. Chesson, P. (2000). Mechanisms of maintenance of species diversity. Annual Review of Ecology and Systematics, 31, 343–366.

10. Cornelissen, J.H.C., Diez, P.C. & Hunt, R. (1996). Seedling growth, allocation and leaf attributes in a wide range of woody plant species and types. Journal of Ecology, 84, 755–765.

11. Cornwell, W.K., Westoby, M., Falster, D.S., FitzJohn, R.G., O’Meara, B.C., Pennell, M.W., McGlinn, D.J., Eastman, J.M., Moles, A.T., Reich, P.B., Tank, D.C., Wright, I.J., Aarssen, L., Beaulieu, J.M., Kooyman, R.M., Leishman, M.R., Miller, E.T., Niinemets, U., Oleksyn, J., Ordonez, A., Royer, D.L., Smith, S.A., Stevens, P.F., Warman, L., Wilf, P. & Zanne, A.E. (2014). Functional distinctiveness of major plant lineages. Journal of Ecology, 102, 345–356.

12. Craine, J.M. & Reich, P.B. (2005). Leaf-level light compensation points in shade-tolerant woody seedlings. New Phytologist, 166, 710–713.

13. De Kauwe, M.G., Medlyn, B.E., Zaehle, S., Walker, A.P., Dietze, M.C., Wang, Y.P., Luo, Y., Jain, A.K., El-Masri, B., Hickler, T., Warlind, D., Weng, E., Parton, W.J., Thornton, P.E., Wang, S., Prentice, I.C., Asao, S., Smith, B., McCarthy, H.R., Iversen, C.M., Hanson, P.J., Warren, J.M., Oren, R. & Norby, R.J. (2014). Where does the carbon go? A model-data intercomparison of vegetation carbon allocation and turnover processes at two temperate forest free-air CO_2_ enrichment sites. New Phytologist, 203, 883–899.

14. Díaz, S., Kattge, J., Cornelissen, J.H.C., Wright, I.J., Lavorel, S., Dray, S., Reu, B., Kleyer, M., Wirth, C., Prentice, I.C., Garnier, E., Bönisch, G., Westoby, M., Poorter, H., Reich, P.B., Moles, A.T., Dickie, J., Gillison, A.N., Zanne, A.E., Chave, J., Wright, S.J., Sheremetev, S.N., Jactel, H., Baraloto, C., Cerabolini, B., Pierce, S., Shipley, B., Kirkup, D., Casanoves, F., Joswig, J.S., Gnther, A., Falczuk, V., Rger, N., Mahecha, M.D. & Gorné, L.D. (2016). The global spectrum of plant form and function. Nature, 529, 167–171.

15. Duursma, R.A. & Falster, D.S. (2016). Leaf mass per area, not total leaf area, drives differences in above-ground biomass distribution among woody plant functional types. New Phytologist, p. Accepted 28/04/2016.

16. Eddelbuettel, D. (2013). Seamless R and C++ integration with Rcpp. Springer, New York.

17. Eddelbuettel, D., Emerson, J.W. & Kane, M.J. (2015). BH: Boost C++ Header Files.

18. Enquist, B.J., Kerkhoff, A.J., Stark, S.C., Swenson, N.G., McCarthy, M.C. & Price, C.A. (2007). A general integrative model for scaling plant growth, carbon flux, and functional trait spectra. Nature, 449, 218–222.

19. Enquist, B.J., West, G.B., Charnov, E.L. & Brown, J.H. (1999). Allometric scaling of production and life-history variation in vascular plants. Nature, 401, 907–911.

20. Falster, D.S., Brännström, Å., Dieckmann, U. & Westoby, M. (2011). Influence of four major plant traits on average height, leaf-area cover, net primary productivity, and biomass density in single-species forests: a theoretical investigation. Journal of Ecology, 99, 148–164.

21. Falster, D.S., Duursma, R.A., Ishihara, M.I., Barneche, D.R., FitzJohn, R.G., Vårhammar, A., Aiba, M., Ando, M., Anten, N., Aspinwall, M.J., Baltzer, J.L., Baraloto, C., Battaglia, M., Battles, J.J., Bond-Lamberty, B., van Breugel, M., Camac, J., Claveau, Y., Coll, L., Dannoura, M., Delagrange, S., Domec, J.C., Fatemi, F., Feng, W., Gargaglione, V., Goto, Y., Hagihara, A., Hall, J.S., Hamilton, S., Harja, D., Hiura, T., Holdaway, R., Hutley, L.S., Ichie, T., Jokela, E.J., Kantola, A., Kelly, J.W.G., Kenzo, T., King, D., Kloeppel, B.D., Kohyama, T., Komiyama, A., Laclau, J.P., Lusk, C.H., Maguire, D.A., le Maire, G., ä, A.M., Markesteijn, L., Marshall, J., McCulloh, K., Miyata, I., Mokany, K., Mori, S., Myster, R.W., Nagano, M., Naidu, S.L., Nouvellon, Y., O’Grady, A.P., O’Hara, K.L., Ohtsuka, T., Osada, N., Osunkoya, O.O., Peri, P.L., Petritan, A.M., Poorter, L., Portsmuth, A., Potvin, C., Ransijn, J., Reid, D., Ribeiro, S.C., Roberts, S.D., Rodriguez, R., Saldana-Acosta, A., Santa-Regina, I., Sasa, K., Selaya, N.G., Sillett, S.C., Sterck, F., Takagi, K., Tange, T., Tanouchi, H., Tissue, D., Umehara, T., Utsugi, H., Vadeboncoeur, M.A., Valladares, F., Vanninen, P., Wang, J.R., Wenk, E., Williams, R., de Aquino Ximenes, F., Yamaba, A., Yamada, T., Yamakura, T., Yanai, R.D. & York, R.A. (2015). BAAD: a Biomass And Allometry Database for woody plants. Ecology, 96, 1445.

22. Falster, D.S., FitzJohn, R.G., Brännström, Å., Dieckmann, U. & Westoby, M. (2016). plant: A package for modelling forest trait ecology and evolution. Methods in Ecology and Evolution, 7, 136–146.

23. Friend, A.D., Lucht, W., Rademacher, T.T., Keribin, R., Betts, R., Cadule, P., Ciais, P., Clark, D.B., Dankers, R., Falloon, P.D., Ito, A., Kahana, R., Kleidon, A., Lomas, M.R., Nishina, K., Ostberg, S., Pavlick, R., Peylin, P., Schaphoff, S., Vuichard, N., Warszawski, L., Wiltshire, A. & Woodward, F.I. (2014). Carbon residence time dominates uncertainty in terrestrial vegetation responses to future climate and atmospheric CO_2_. Proceedings of the National Academy of Sciences, 111, 3280–3285.

24. Gibert, A., Gray, E.F., Westoby, M., Wright, I.J. & Falster, D.S. (2016). On the link between functional traits and growth rate: meta-analysis shows effects change with plant size, as predicted. Journal of Ecology, 104, 1488–1503.

25. Givnish, T.J. (1988). Adaptation to sun and shade: a whole-plant perspective. Australian Journal of Plant Physiology, 15, 63–92.

26. Gower, S.T., Reich, P.B. & Son, Y. (1993). Canopy dynamics and aboveground production of five tree species with different leaf longevities. Tree Physiology, 12, 327–345.

27. Grime, J.P. (1977). Evidence for the existence of three primary strategies in plants and its relevance to ecological and evolutionary theory. American Naturalist, 111, 1169.

28. Hérault, B., Bachelot, B., Poorter, L., Rossi, V., Bongers, F., Chave, J., Paine, C.E.T., Wagner, F. & Baraloto, C. (2011). Functional traits shape ontogenetic growth trajectories of rain forest tree species. Journal of Ecology, 99, 1431–1440.

29. Huxley, J.S. (1932). Problems of Relative Growth. Methuen & Co. Ltd.

30. Iida, Y., Kohyama, T.S., Swenson, N.G., Su, S.H., Chen, C.T., Chiang, J.M. & Sun, I.F. (2014). Linking functional traits and demographic rates in a subtropical tree community: the importance of size dependency. Journal of Ecology, pp. 641–650.

31. King, D.A. (1999). Juvenile foliage and the scaling of tree proportions, with emphasis on eucalyptus. Ecology, 80, 1944–1954.

32. King, D.A. (2005). Linking tree form, allocation and growth with an allometrically explicit model. Ecological Modelling, 185, 77–91.

33. King, D.A. (2011). Size-related changes in tree proportions and their potential influence on the course of height growth. In: Size- and Age-Related Changes in Tree Structure and Function (eds. Meinzer, F.C.C., Lachenbruch, B., Dawson, T.E.E., Meinzer, F.C. & Niinemets, U.). Springer Netherlands, vol. 4 of Tree Physiology, pp. 165–191.

34. Kneeshaw, D.D., Kobe, R.K., Coates, K.D. & Messier, C. (2006). Sapling size influences shade tolerance ranking among southern boreal tree species. Journal of Ecology, 94, 471–480.

35. Koch, G.W., Sillett, S.C., Jennings, G.M. & Davis, S.D. (2004). The limits to tree height. Nature, 428, 851–854.

36. Kokko, H. (2007). Modelling for field biologists and other interesting people. Cambridge University Press, Cambridge.

37. Kunstler, G., Falster, D., Coomes, D.A., Hui, F., Kooyman, R.M., Laughlin, D.C., Poorter, L., Vanderwel, M., Vieilledent, G., Wright, S.J., Aiba, M., Baraloto, C., Caspersen, J., Cornelissen, J.H.C., Gourlet-Fleury, S., Hanewinkel, M., Herault, B., Kattge, J., Kurokawa, H., Onoda, Y., Peñuelas, J., Poorter, H., Uriarte, M., Richardson, S., Ruiz-Benito, P., Sun, I.F., Ståhl, G., Swenson, N.G., Thompson, J., Westerlund, B., Wirth, C., Zavala, M.A., Zeng, H., Zimmerman, J.K., Zimmermann, N.E. & Westoby, M. (2016). Plant functional traits have globally consistent effects on competition. Nature, 529, 204–207.

38. Lambers, H. & Poorter, H. (1992). Inherent variation in growth rate between higher plants: A search for physiological causes and ecological consequences. In: Advances in Ecological Research (ed. M. Begon and A.H. Fitter). Academic Press, vol. 23, pp. 187–261.

39. Li, G., Harrison, S.P., Prentice, I.C. & Falster, D.S. (2014). Simulation of tree-ring widths with a model for primary production, carbon allocation, and growth. Biogeosciences, 11, 6711–6724.

40. Lusk, C.H., Falster, D.S., Jara-Vergara, C.K., Jimenez-Castillo, M. & Saldaña-Mendoza, A. (2008). Ontogenetic variation in light requirements of juvenile rainforest evergreens. Functional Ecology, 22, 454–459.

41. Lusk, C.H. & Jorgensen, M.A. (2013). The whole-plant compensation point as a measure of juvenile tree light requirements. Functional Ecology, 27, 1286–1294.

42. Mäkelä, A. (1997). A carbon balance model of growth and self-pruning in trees based on structural relationships. Forest Science, 43, 7–24.

43. Messier, C., Doucet, R., Ruel, J.C., Claveau, Y., Kelly, C. & Lechowicz, M.J. (1999). Functional ecology of advance regeneration in relation to light in boreal forests. Canadian Journal of Forest Research, 29, 812–823.

44. Moles, A.T. & Westoby, M. (2006). Seed size and plant strategy across the whole life cycle. Oikos, 113, 91–105.

45. Moorcroft, P.R., Hurtt, G.C. & Pacala, S.W. (2001). A method for scaling vegetation dynamics: the ecosystem demography model (ED). Ecological Monographs, 71, 557–586.

46. Niinemets, Ü. (2010). A review of light interception in plant stands from leaf to canopy in different plant functional types and in species with varying shade tolerance. Ecological Research, 25, 693–714.

47. Osunkoya, O.O. (1996). Light requirements for regeneration in tropical forest plants: Taxon-level and ecological attribute effects. Australian Journal of Ecology, 21, 429–441.

48. Paine, C.E.T., Amissah, L., Auge, H., Baraloto, C., Baruffol, M., Bourland, N., Bruelheide, H., Danou, K., de Gouvenain, R.C., Doucet, J.L., Doust, S., Fine, P.V.A., Fortunel, C., Haase, J., Holl, K.D., Jactel, H., Li, X., Kitajima, K., Koricheva, J., Martínez-Garza, C., Messier, C., Paquette, A., Philipson, C., Piotto, D., Poorter, L., Posada, J.M., Potvin, C., Rainio, K., Russo, S.E., Ruiz-Jaen, M., Scherer-Lorenzen, M., Webb, C.O., Wright, S.J., Zahawi, R.A. & Hector, A. (2015). Globally, functional traits are weak predictors of juvenile tree growth, and we do not know why. Journal of Ecology, 103, 978–989.

49. Pérez-Harguindeguy, N., Díaz, S., Garnier, E., Lavorel, S., Poorter, H., Jaureguiberry, P., Bret-Harte, M.S., Cornwell, W.K., Craine, J.M., Gurvich, D.E., Urcelay, C., Veneklaas, E.J., Reich, P.B., Poorter, L., Wright, I.J., Ray, P., Enrico, L., Pausas, J.G., de Vos, A.C., Buchmann, N., Funes, G., Quétier, F., Hodgson, J.G., Thompson, K., Morgan, H.D., ter Steege, H., van der Heijden, M.G.A., Sack, L., Blonder, B., Poschlod, P., Vaieretti, M.V., Conti, G., Staver, A.C., Aquino, S. & Cornelissen, J.H.C. (2013). New handbook for standardised measurement of plant functional traits worldwide. Australian Journal of Botany, 61, 167–234.

50. Poorter, L. & Bongers, F. (2006). Leaf traits are good predictors of plant performance across 53 rain forest species. Ecology, 87, 1733–1743.

51. Poorter, L., Wright, S.J., Paz, H., Ackerly, D., Condit, R., Ibarra-Manriques, G., Harms, K., Licona, J., Martinez-Ramos, M., Mazer, S., Muller-Landau, H.C., Pena-Claros, M., Webb, C. & Wright, I. (2008). Are functional traits good predictors of demographic rates? Evidence from five Neotropical forests. Ecology, 89, 1908–1920.

52. Purves, D. & Pacala, S. (2008). Predictive models of forest dynamics. Science, 320, 1452–1453.

53. R Core Team (2015). R: a language and environment for statistical computing. R Foundation for Statistical Computing, Vienna, Austria.

54. Rees, M., Condit, R., Crawley, M., Pacala, S.W. & Tilman, D. (2001). Long-term studies of vegetation dynamics. Science, 293, 650–655.

55. Rees, M., Osborne, C.P., Woodward, F.I., Hulme, S.P., Turnbull, L.A. & Taylor, S.H. (2010). Partitioning the components of relative growth rate: how important is plant size variation? The American Naturalist, 176, E152–E161.

56. Reich, P.B., Walters, M.B. & Ellsworth, D.S. (1992). Leaf life-span in relation to leaf, plant, and stand characteristics among diverse ecosystems. Ecological Monographs, 62, 365–392.

57. Rüger, N., Wirth, C., Wright, S.J. & Condit, R. (2012). Functional traits explain plasticity of growth rates in tropical tree species. Ecology, 93, 2626–2636.

58. Ryan, M.G., Phillips, N. & Bond, B.J. (2006). The hydraulic limitation hypothesis revisited. Plant, Cell & Environment, 29, 367–381.

59. Schäling, B. (2014). The Boost C++ libraries. 2nd edn. XML Press.

60. Shinozaki, K., Yoda, K., Hozumi, K. & Kira, T. (1964). A quantitative analysis of plant form - the pipe model theory. I. Basic analyses. Japanese Journal of Ecology, 14, 97–105.

61. Sillett, S.C., Van Pelt, R., Koch, G.W., Ambrose, A.R., Carroll, A.L., Antoine, M.E. & Mifsud, B.M. (2010). Increasing wood production through old age in tall trees. Forest Ecology and Management, 259, 976–994.

62. Sitch, S., Huntingford, C., Gedney, N., Levy, P.E., Lomas, M., Piao, S.L., Betts, R., Ciais, P., Cox, P., Friedlingstein, P., Jones, C.D., Prentice, I.C. & Woodward, F.I. (2008). Evaluation of the terrestrial carbon cycle, future plant geography and climate-carbon cycle feedbacks using five Dynamic Global Vegetation Models (DGVMs). Global Change Biology, 14, 2015–2039.

63. Stephenson, N.L., Das, A.J., Condit, R., Russo, S.E., Baker, P.J., Beckman, N.G., Coomes, D.A., Lines, E.R., Morris, W.K., Rüger, N., Ivarez, E., Blundo, C., Bunyavejchewin, S., Chuyong, G., Davies, S.J., Duque,., Ewango, C.N., Flores, O., Franklin, J.F., Grau, H.R., Hao, Z., Harmon, M.E., Hubbell, S.P., Kenfack, D., Lin, Y., Makana, J.R., Malizia, A., Malizia, L.R., Pabst, R.J., Pongpattananurak, N., Su, S.H., Sun, I.F., Tan, S., Thomas, D., van Mantgem, P.J., Wang, X., Wiser, S.K. & Zavala, M.A. (2014). Rate of tree carbon accumulation increases continuously with tree size. Nature, 507, 90–93.

64. Thomas, S.C. (2011). Age-related changes in tree growth and functional biology: the role of reproduction. In: Size- and Age-Related Changes in Tree Structure and Function (eds. Meinzer, F.C., Lachenbruch, B. & Dawson, T.E.). Springer Netherlands, Dordrecht, vol. 4, pp. 33–64.

65. Thomas, S.C. & Bazzaz, F.A. (1999). Asymptotic height as a predictor of photosynthetic characteristics in malaysian rain forest trees. Ecology, 80, 1607–1622.

66. Thornley, J.H.M. & Cannell, M.G.R. (2000). Modelling the components of plant respiration: representation and realism. Annals of Botany, 85, 55–67.

67. Van Kleunen, M., Weber, E. & Fischer, M. (2010). A meta-analysis of trait differences between invasive and non-invasive plant species. Ecology Letters, 13, 235–245.

68. Visser, M.D., Bruijning, M., Wright, S.J., Muller-Landau, H.C., Jongejans, E., Comita, L.S. & de Kroon, H. (2016). Functional traits as predictors of vital rates across the life cycle of tropical trees. Functional Ecology, 30, 168–180.

69. Warton, D.I., Wright, I.J., Falster, D.S. & Westoby, M. (2006). Bivariate line-fitting methods for allometry. Biological Reviews, 81, 259–291.

70. Wenk, E.H. & Falster, D.S. (2015). Quantifying and understanding reproductive allocation schedules in plants. Ecology and Evolution, 5, 5521–5538.

71. Westoby, M., Falster, D.S., Moles, A.T., Vesk, P. & Wright, I.J. (2002). Plant ecological strategies: some leading dimensions of variation between species. Annual Review of Ecology and Systematics, 33, 125–159.

72. Wilson, G., Aruliah, D.A., Brown, C.T., Hong, N.P.C., Davis, M., Guy, R.T., Haddock, S.H.D., Huff, K., Mitchell, I.M., Plumbley, M., Waugh, B., White, E.P. & Wilson, P. (2014). Best practices for scientific computing. PLoS Biology, 12, e1001745.

73. Wright, I.J., Reich, P.B., Westoby, M., Ackerly, D., Baruch, Z., Bongers, F., Cavender-Bares, J., Chapin, F., Cornelissen, J., Diemer, M., Flexas, J., Garnier, E., Groom, P., Gulias, J., Hikosaka, K., Lamont, B., Lee, T., Lee, W., Lusk, C., Midgley, J., Navas, M.L., Niinemets, Ű., Oleksyn, J., Osada, N., Poorter, H., Poot, P., Prior, L., Pyankov, V., Roumet, C., Thomas, S., Tjoelker, M., Veneklaas, E. & Villar, R. (2004). The world-wide leaf economics spectrum. Nature, 428, 821–827.

74. Wright, I.J. & Westoby, M. (2000). Cross-species relationships between seedling relative growth rate, nitrogen productivity and root vs leaf function in 28 australian woody species. Functional Ecology, 14, 97–107.

75. Wright, S.J., Kitajima, K., Kraft, N.J.B., Reich, P.B., Wright, I.J., Bunker, D.E., Condit, R., Dalling, J.W., Davies, S.J., Díaz, S., Engelbrecht, B.M.J., Harms, K.E., Hubbell, S.P., Marks, C.O., Ruiz-Jaen, M.C., Salvador, C.M. & Zanne, A.E. (2010). Functional traits and the growth-mortality trade-off in tropical trees. Ecology, 91, 3664–3674.

76. Yokozawa, M. & Hara, T. (1995). Foliage profile, size structure and stem diameter plant height relationship in crowded plant-populations. Annals of Botany, 76, 271–285.

77. Zanne, A.E., Westoby, M., Falster, D.S., Ackerly, D.D., Loarie, S.R., Arnold, S.E. & Coomes, D.A. (2010). Angiosperm wood structure: global patterns in vessel anatomy and their relation to wood density and potential conductivity. American Journal of Botany, 97, 207–215.

